# Targeting mTOR restores tau-induced metabolic, mitochondrial, and cognitive deficits in a tauopathy mouse model

**DOI:** 10.1101/2024.11.24.625068

**Authors:** Zhi Tang, Min Guo, Yuanting Ding, Yi Wen, Bo Li, Yan Xiao, Ruiqing Ni, Zhizhong Guan, Xiao-lan Qi

## Abstract

**Aim:** Hyperphosphorylated tau is a hallmark of Alzheimer’s disease (AD). However, whether mammalian target of rapamycin (mTOR) directly regulates tau phosphorylation at Ser214, Ser356, and Thr231 and how such regulation affects mitochondrial function remain unclear. In this study, we aimed to determine whether mTOR-mediated tau phosphorylation disrupts mitochondrial and metabolic profiles and cognitive function and whether these effects can be reversed by the mTOR inhibitor rapamycin.

**Methods:** Tau^3E^-overexpressing mice were generated by bilateral injection of adeno-associated virus vectors encoding the phosphomimetic TauS214E/T231E/S356E (Tau^3E^) variant into the hippocampal CA3 region of 2-month-old C57BL/6 mice. The mice received intraperitoneal rapamycin for one week. Cognitive performance was assessed using the Morris water maze. MALDI-mass spectrometry imaging was conducted to evaluate metabolic alterations in the brain tissue slices. mTOR, p70S6K, and tau phosphorylation protein expression; mitochondrial dynamics markers; and reactive oxygen species (ROS) levels were analyzed by Western blotting, immunofluorescence, and flow cytometry in HT22 cells, Tau^3E^ mice, and postmortem AD brain tissues.

**Results:** Phosphorylated mTORS2448 colocalized with p-TauSer214, p-TauSer356, and p-TauThr231 in the hippocampal CA3 region of AD brains. HT22 cells and Tau^3E^ mice exhibited increased levels of p-mTOR, p-p70S6K, and ROS production; mitochondrial fragmentation; and tau phosphorylation at Ser214, Ser356, and Thr231. Rapamycin treatment partially ameliorated cognitive deficits, reduced oxidative stress and mitochondrial dysfunction, and restored metabolic profile particularly the inosine-pentose phosphate pathway-glycolysis metabolic axis, structural lipidome axis and the neurotransmitter homeostasis.

**Conclusion:** mTOR activation contributes to site-specific tau hyperphosphorylation, mitochondrial dysfunction, and cognitive impairment. Pharmacological inhibition of mTOR by rapamycin attenuated tau pathology, regulated metabolic alterations, preserved mitochondrial homeostasis, and improved cognitive function, suggesting a potential therapeutic strategy for AD.

## Introduction

Alzheimer’s disease (AD) is the most common neurodegenerative disease characterized by progressive cognitive decline. Extracellular amyloid-β (Aβ) plaques and neurofibrillary tangles (NFTs) formed by the hyperphosphorylation of the microtubule-associated protein tau are important hallmarks of AD (*1*). In AD, tau protein is found mainly in neurons, whereas in corticobasal degeneration and progressive supranuclear palsy, tau is located mainly in astrocytes (*2, 3*). Tau plays important cellular and physiological roles and regulates the assembly, transport and stability of microtubules and neuronal synaptic function (*4*). Mutations in tau coding regions, disruptions in tau mRNA splicing, alterations in tau posttranslational modifications, and cellular stress factors such as oxidative stress and inflammation all contribute to an increased tendency for tau to aggregate and hinder its clearance (*4*). In pathological states, tau aggregates are internalized by adjacent cells through endocytosis, pinocytosis, and phagocytosis; are transferred from one cell to another; and spread in different regions of the brain in a prion-like manner (*5*). Tau aggregate accumulation is associated with cognitive decline (*6*) and leads to neuroinflammation, atrophy, and metabolic and microstructural changes (*7–13*). Recent studies have shown that the combination of brain glucose hypometabolism and hyperphosphorylated Tau (p-Tau) pathology is a clinical predictor of imminent cognitive decline (*14–16*).

Tau contains 85 potentially phosphorylated (p-) serine (Ser), threonine (Thr) and tyrosine (Tyr) residues in its longest isoform (2N4R)(*17*). Hyperphosphorylation of tau can cause microtubule depolymerization, blockage of axoplasmic flow and neuronal cross-linking, resulting in neuronal degeneration and loss (*17*). Mass spectrometry and specific antibody immunostaining revealed 10 phosphorylation sites in soluble tau isolated from normal human brains (*18*), and approximately 45 different serine, threonine and tyrosine phosphorylation sites were found in insoluble tau isolated from the brains of AD patients (*18*). Tau phosphorylation is a clinical biomarker and therapeutic target in AD and other tauopathies (*17, 19*). p-Thr181, p-Thr217 (*20*), and p-Thr231 (*21*) are increased in the cerebrospinal fluid (CSF) of AD patients and have been used as diagnostic biomarkers. Moreover, the level of CSF p-Thr231 is particularly correlated with the cortical tangle load in AD patients. These phosphorylation sites affect the ability of tau to bind to microtubules, which is associated with mitochondrial dysfunction and homeostatic imbalance in neurons (*22*).

Mammalian target of rapamycin (mTOR) is a serine/threonine protein kinase that regulates the basic bioprocessing of cells and is known to phosphorylate tau to facilitate its assembly into NFT, and the “mTOR signaling pathway” plays an important role in AD and aging (*23*). mTOR controls many cellular functions, such as cell growth, proliferation, differentiation, survival, autophagy, and metabolism (*24, 25*). The formation and aggregation of NFT and Ab plaques are significantly regulated by mTOR signaling (*26*). Several clinical trials assessing the efficacy of rapamycin, an mTOR inhibitor for AD, are ongoing (*27*), including a phase 2 trial (REACH NCT04629495) in patients with mild cognitive impairment and early-stage AD (*28*). We previously revealed that mTOR directly phosphorylates tau at Ser214, Ser356 and Thr231 in cell lines. mTOR regulates tau production and aggregation and affects the activity of protein kinases, such as protein kinases A and B and glycogen synthase kinase-3, resulting in compromised microtubule stability and intracellular tau accumulation and translocation (*26, 29–32*). However, whether mTOR directly interacts with tau at Ser214, Ser356 and Thr231 in AD animal models and in the brains of AD patients is not known.

In the present study, we aimed to establish causal links among tau phosphorylation, mTOR, reactive oxygen species (ROS), mitochondrial dysfunction and cognitive impairment. We hypothesized that 1) mTOR-regulated tau phosphorylation at Ser214/Ser356/Thr231 affects mitochondrial homeostasis and neuronal damage in an AD model and that 2) rapamycin can ameliorate p-Tau^S214E/T231E/S356E^ (Tau^3E^)-induced cognitive impairment, mitochondrial dysfunction and ROS (*23, 33, 34*). We evaluated p-tau levels, mTOR levels, mitochondrial dynamics, and ROS levels in AD postmortem brain tissues, HT22 cells and Tau^3E-^overexpressing mice. We assessed the effects of treatment with rapamycin (i.p.) in Tau^3E-^overexpressing mice on the cognition, molecular and metabolomic profiles by using MALDI- mass spectrometry imaging (MSI).

## 2 Materials and methods

### 2.1 Materials and antibodies, plasmids, and viruses

Rapamycin, tris aminomethane (Tris), radioimmunoprecipitation assay (RIPA), sodium dodecyl sulfate (SDS) buffer and protease inhibitor cocktail were obtained from Sigma Aldrich Co. (St, Louis, MO, USA). A Bradford kit was purchased from Bio-Rad (California, USA). For detailed information on the primary antibodies and reagents used in the present study, please refer to **STables 1 and 2**.

Construction of the recombinant Tau protein vector: Using a full-length human Tau40 (a gift from Prof. Jianzhi Wang, Huazhong University of Science and Technology, China) as a template, we used overlap extension polymerase chain reaction (PCR) to construct a point mutation and converted it into the p-Tau protein model: glutamate (Glu: GAA) TauS214E/T231E/S356E and point mutations to the dephosphorylated Tau protein model: alanine (Ala: GCA) TauS214A, TauT231A, TauS356A and TauS214A/T231A/S356A, which were seamlessly connected by restriction enzyme digestion into the expression vector p-enhanced green fluorescent protein (EGFP)-C1, which has been optimized for brighter fluorescence and higher expression in mammalian cells: 1) The gene sequence of full-length human Tau40 was obtained from the NCB database (ID: 4137). 2) Primer design: For the Ser214, Thr231, and Ser356 positions in the MAPT amino acid sequence, mutation primers were designed via GenTool V_8.9 software, and the corresponding subcloning enzyme cleavage sites were added (BgIII and EcoRI). The primers used are listed in **STables 3** and **4**. 3) Overlap PCR amplification: Overlap extension PCR involves the design of 2 pairs of primers: the Tau40-Forward and Tau40-Reverse primers are at the ends of the tau gene, whereas the other primers are in the middle of the tau gene. For example, the overlap extension PCR procedure is as follows: First, we used two pairs of primers, namely, the primers Tau40-Forward/TauS214E, TauT231E, and TauS356E-Forward and the primers Tau40-Reverse/TauS214E, TauT231E, and TauS356E-Reverse, to generate two DNA fragments using the target gene as the template. The PCR products were gel-purified. These two DNA fragments were therefore able to anneal together with their complementary overhanging cohesive ends, which were then extended and amplified via PCR via the primer pair Tau40-Forward/Tau40-Reverse to obtain a full-length gene with the expected mutation. After the end of the reaction, the results were analyzed via agarose gel electrophoresis. 4) Double enzyme digestion was performed according to the electrophoresis results of the PCR products, the products were recovered from the gel, and the expression vector pEGFP-C1 was digested with BgIII and EcoRI endonucleases via the following enzyme digestion system: reaction conditions: 37 °C, 6 h; after enzyme digestion, the results were examined by agarose gel electrophoresis. 5) The product of empty pEGFP-C1 and the recovery of the tau protein mutation were ligated with T4 ligase. After sample addition, ligation was performed at 4 °C overnight. The ligation products were transformed into DH5α coli cells, and positive clones were sequenced (Sangon Bioengineering Co., Ltd., Shanghai, China) to ensure the success of the construction of the recombinant plasmid.

### 2.2 Postmortem human brain tissue

Five AD patients, each with a clinical diagnosis confirmed by pathological examination of amyloid and tau, were included in this study (detailed information in **STable 5**) (*35, 36*). Paraffin-embedded autopsied hippocampal tissues were obtained from the Netherlands Brain Bank (NBB), Netherlands. The study was conducted in accordance with the principles of the Declaration of Helsinki and subsequent revisions. All the experiments on autopsied human brain tissue were carried out in accordance with ethical approval obtained from the regional human ethics committee of Guiyang Hospital (2022–290) and the medical ethics committee of the VU Medical Center for NBB tissue.

### 2.3 Cell culture and transfection

The cell culture mixture was prepared as described previously (*37, 38*). Mouse hippocampal HT22 cells were grown to 70-80% confluence in 75 cm^2^ plastic culture flasks (Corning, China) in a mixture of 5% CO_2_ and 95% air at 37 °C, and Dulbecco’s modified Eagle’s medium /F12 medium (1:1) supplemented with 10% fetal bovine serum, 100 units/mL penicillin, and 100 mg/mL streptomycin was used. Different plasmids expressing either h-Tau (Tau40) or tau mutants were transfected into HT22 cells via Lipofectamine™ 2000 (Invitrogen, USA). Briefly, the cells were split into 6-well plates at 60% confluency. For each well, 2 μg of DNA was transfected with Lipofectamine™ 2000 according to the manufacturer’s instructions. The cells were harvested at 48 h after transfection.

### 2.4 Protein extraction and western blotting

HT22 cells and the hippocampi of the mice were lysed in RIPA buffer supplemented with 0.1% protease inhibitor cocktail on ice. The protein concentration was measured with a Bradford kit (Bio-Rad, USA). The proteins were analyzed via western blotting as previously described (*39*). The lysates were separated on 4–12% sodium dodecyl sulfate–polyacrylamide gels (Absin, China), and the proteins were transferred onto 0.22/0.45 μm polyvinylidene difluoride membranes. After the membranes were blocked with 5% milk, they were incubated with primary antibodies (**STable 1**) at 4 °C overnight and then incubated with secondary peroxidase-coupled anti-mouse or anti-rabbit antibodies (1:5000) at room temperature for 2 h. The immunoreactive bands were visualized with Immobilon Western Horseradish peroxidase substrate luminol reagent (Millipore, USA) via a ChemiDoc™ MP imaging system (Bio-Rad, USA).

### 2.5 ROS detection

ROS levels were detected as described previously (*32, 40, 41*). Briefly, HT22 cells were seeded at 1×10^5^ cells per well in 6-well plates and transfected with the Tau^3E^ plasmid at 48 h. These cells were subsequently incubated with 5 μM dihydroethidium (DHE) for 30 min at 37 °C. HT22 cells were rotated gently, washed and resuspended in PBS. The mean fluorescence intensity was detected by a NovoCyte flow cytometer (Agilent Technologies, USA). The data were analyzed via NovoExpress software.

### 2.6 Animal surgery and treatment

Forty wild-type C57BL/6 mice (2 months old, male) were included in the study (Guizhou Experimental Animal Center). All the mice were housed in ventilated cages at a controlled temperature of 22 ± 2 °C, humidity of 50 ± 5%, and a circadian rhythm of 12:12 h. Poplar wood shavings were placed in the cages for environmental enrichment. Food (safe and sterilized) and water (softened and sterilized) were provided ad libitum. All experimental protocols were approved by the Guiyang Regional Animal Care Center and Ethics Committee. The mice were randomly assigned to four groups (the control group, vector group, Tau^3E^ group and Tau^3E^ +rapamycin group; n = 10/group). The mice were deeply anesthetized with an initial dose of 5% isoflurane in an oxygen/air mixture (1:4, 1 L/min) and maintained at 1.5% isoflurane in an oxygen/air mixture (1:4, 0.6 L/min). Anesthetized mice were placed on a stereotaxic apparatus (RWD Life Science, China), and the coordinates for injection were 2.2 mm posterior, 2.6 mm lateral, and 2.3 mm ventral to the bregma. Using a microinjection system (RWD Life Science, Shenzhen, China), AAV-hSyn-EGFP-Tau^S214E/T231E/S356E^ or the control vector (4.25×10^12^ viral genome/ml) was injected into the bilateral hippocampal CA3 region at a rate of 0.125 μl/min, while the control group was given an equal dose of sterile 0.9% saline, and the needle was maintained for 10 min before withdrawal. The body temperature and respiratory rate of the mice were monitored during surgery. The body temperature of each animal was maintained at 36.5 ± 0.5 °C throughout the procedure via a warming pad. Lidocaine ointment was wiped locally to the scalp to reduce pain. Four weeks after virus infusion, the mice in the Tau^3E^ + rapamycin group were administered rapamycin (1.5 mg/kg body weight) via intraperitoneal (i.p.) injection three times for one week on alternate days. Behavioral tests were subsequently performed after the rapamycin treatment period.

### 2.7 Behavioral testing

The Morris water maze test was performed from 2:00 p.m. to 8:00 p.m. as described previously (*38*). During Morris water maze training, all the mice were subjected to 4 training trials daily for four consecutive days. In each trial, the mice were trained to find a 20 cm diameter hidden platform submerged 1 cm under the water surface for 60 s. Afterwards, the mice were kept on the platform for 10 s. If these mice could not find the platform within 60 s, they were guided onto the platform within 60 s and kept on the platform for 20 s afterwards. On day 5, a spatial probe trial was executed in which the platform was removed. The dependent variables used for the analysis were latency (seconds) to reach the platform for the learning trials and the number of platform crossings, latency to first cross the platform location (seconds), and velocity (cm/second) for the probe trial. All trials were video recorded, and the data were analyzed via the EthoVisionXT system (Noldus Information Technology). After the behavioral tests were performed, all the mice were sacrificed under deep anesthesia with pentobarbital sodium (50 mg/kg body weight) and transcardiacly perfused with phosphate-buffered saline (PBS, pH 7.4). The left hemisphere of the mouse brain was stored at -80 °C and used for western blotting. The right hemisphere of the mouse brain was fixed in 4% paraformaldehyde in 1 × PBS (pH 7.4) for 24 h and stored in 1 × PBS (pH 7.4) at 4 °C. For immunofluorescence staining, the fixed right hemisphere brain tissues were dehydrated via a vacuum infiltration processor (Leica ASP200S, Germany) and embedded in paraffin via an Arcadia H heated embedding workstation (Leica, Germany).

### 2.7 Immunofluorescence staining and confocal imaging

Paraffin-embedded hippocampal tissue blocks from AD patients were cut into 6 µm pieces via a microtome (HM340E; Thermo Fisher Scientific, USA). The tissue sections were deparaffinized and rehydrated prior to antigen retrieval (citrate buffer, pH 6.0) in a microwave for 15 min at 98 °C (*7, 42*). These sections were blocked in Tris-buffered saline (TBS) with Tween-20 with 5% goat serum for 60 min. Then, the sections were incubated with primary antibodies (**STable 1**) at 4 °C overnight. Immunoreactions were detected using Alexa Fluor 488- or 546-conjugated secondary antibodies.

To image the cells, after transfection, the HT22 cells were placed on coverslips, rinsed with PBS, and then fixed in 4% paraformaldehyde for 30 min. The cells were permeabilized in 0.1% Triton X-100 in TBS for 10 min. The nonspecific binding sites were blocked with blocking solution (5% bovine serum albumin, 0.1% Triton X-100 in TBS) for 2 h. The cells were incubated with primary antibodies (**STable** 1), anti-4-hydroxynonenal (4-HNE), anti-8-hydroxy-2’-deoxyguanosine (8-OHdG), anti-nicotinamide adenine dinucleotide phosphate oxidase 4 (NOX4) and neuronal nuclei (NeuN), at 4 °C overnight. After the samples were washed with TBS, immunoreactions were detected via Alexa Fluor 488-IgGs or Alexa Fluor 546-IgGs (1:200 for both, Invitrogen, USA). Anti-fading mounting medium supplemented with 4′,6-diamidino-2-phenylindole (DAPI) (Vector Laboratories, USA) was used. The imaging of tau expression was performed at 488 nm for the EGFP-tau (hTau, TauS214A/T231A/S356A (Tau3A), TauS214A, TauT231A, TauS356A, TauS214E, TauT231E, and TauS356E, TauS214 E/T231E/S356E (Tau^3E^)) signals. The sections were mounted with vector anti-fading mounting medium containing DAPI. Images were obtained via a confocal microscope (Olympus, Japan) at 100× magnification for 2 views × 3 slides per group. Confocal images were processed with OlyVIA software (Olympus OlyVIA3.3; Olympus, Japan). The nuclearLJcytoplasmic ratio of EGFP-tau was determined by using ImageJ with thresholding and analysis tools. The mean fluorescence intensities of 4-HNE, 8-OHdG, NOX4 and Tomm20 were quantified via automatic thresholding of the fluorescence intensity via ImageJ software as described previously. Mitochondrial morphology was analyzed in HT22 cells using the following criteria: the mitochondria in normal HT22 cells were filamentous and linearly tubular, whereas the mitochondria in mitochondria-fragmented HT22 cells were shortened and punctate; HT22 cells with more than 70% mitochondrial fragmentation were identified as mitochondrially fragmented. Each experiment was repeated three times, and at least 60 HT22 cells per group were analyzed.

For imaging of the mouse brain, coronal sections of the paraffin-embedded mouse half hemispheres were cut at a thickness of 6 μm via a microtome. NeuN, 4-HNE, 8-OHdG and NOX4 staining was performed as described previously (*7, 38*). The imaging of tau expression (EGFP-Tau^3E^) in the mouse hemi-brain was performed by mounting with anti-fading mounting medium supplemented with DAPI (Vector Laboratories, USA). Overview images of the hemisphere were captured at ×10 magnification using a confocal microscope (Olympus, Japan) and integrated automatically. Images of the sections in the pyramidal layer of the CA3 region were obtained via a confocal microscope (Olympus, Japan) at 100× magnification. Confocal images were processed with OlyVIA software (Olympus OlyVIA3.3; Olympus, Japan). The mean fluorescence intensities of 4-HNE, 8-OHdG and NOX4 were quantified via automatic thresholding of the fluorescence intensity via ImageJ software (NIH, USA). NeuN/DAPI-double-positive cells were counted and analyzed as a percentage of the total number of DAPI-positive cells. The experiments were performed on 6 sections, and a total of 30 cells from each group were analyzed.

### 2.8 Matrix-Assisted Laser Desorption/Ionization (MALDI) matrix preparation and application

The tissues were stored at -80 °C prior to analysis. The tissues were changed to stored at -20 °C for 1 h before use, and tissue sections were prepared using a Leica CM1950 cryostat (Leica Microsystems GmbH, Wetzlar, Germany). Tissue sections (10 μm thick) were thaw mounted onto indium-tin-oxide -coated microscopic slides (Bruker) for MALDI MSI. The mounted tissue sections were dried for 15 minutes in a desiccator prior to matrix application and then transferred to a -80 °C refrigerator for sealing storage after vacuum packaging. *N*-(1-naphthyl) ethylenediamine dihydrochloride (5 mg/mL) was dissolved in 70% HPLC grade acetonitrile with 0.1% trifluoroacetic acid. The 10 μm thick tissue sections were sprayed using a TM-Sprayer (HTX Technologies). The parameters of the matrix application set in the TM-sprayer were as follows: spray nozzle velocity (1200 mm/min), track spacing (3 mm), flow rate (0.08 mL/min), spray nozzle temperature (60 °C), and nitrogen gas pressure (10 pounds per square inch).

### 2.9 Mass spectrometry imaging and data analysis

Metabolites in the samples were imaged using a timsTOF fleX MALDI 2 (Bruker, Germany) equipped with a 10 kHz smartbeam 3D laser. MALDI2 MSIs were performed in positive ion mode in full scan mode for 50–1300 m/z. MALDI-2 MSI was performed at 50 μm spatial resolution. The frequency of the laser was set to 10,000 Hz, with 70% of the laser energy and 200 laser shots per 50 μm pixel.

MSI data were analyzed using SCiLS Lab software (version 2024b premium; Bruker Daltonics, Billerica, MA) with root mean square normalization. Metabolites were identified by comparison of their m/z values (<10 ppm) with those in the Human Metabolome Database. The data were analyzed using the Seurat R package, with the FindMarkers function applying the Wilcoxon test and the Bonferroni correction method to select differentially expressed metabolites. Principal component analysis (PCA) was applied to distinguish the characteristics of metabolite differences between the two groups.

**|**avg_log2FC| > 0.1 and p value < 0.05 were used to screen significantly changed metabolites. Kyoto Encyclopedia of Genes and Genomes (KEGG) enrichment analysis of different metabolites was performed using clusterProfiler 4.9.1.

### 2.10 Statistical analysis

Statistical analysis was performed via GraphPad Prism 8.0 software (GraphPad Software Inc., USA). The results are presented as the mean ± SEMs. The Morris water maze behavior data were assessed via two-way repeated-measures ANOVA. The other parameters were analyzed via one-way ANOVA followed by Tukey’s post hoc test for multiple comparisons. Significance was set at p < 0.05.

## 3 Results

### 3.1 Tau^3E^ increased tau expression and the cellular localization of pTau

The use of the overlap extension PCR point mutagenesis method to perform site-directed mutation of the Tau protein into a phosphorylated model or dephosphorylated Tau has become important for studying the pathological mechanisms of the Tau protein (*43*) (**Fig. 1**). Earlier studies mutated Ser/Thr to Glu to generate pseudo-p-Tau protein and demonstrated that the abnormal hyperphosphorylation of Tau leads to its aggregation, destruction of the microtubule network and cell death. Here, we chose to use the overlapping PCR point mutation method to construct the site-directed phosphorylation vector Tau^3E^ via the full-length human tau protein MAPT and used this method to explore the potential roles of the three sites TauSer214, TauThr231, and TauSer356 (*43*). P-Tau214 and p-Tau231 have a phosphorylation site (Ser214, T231) within the same proline-rich region of tau (**Fig. 1G**). Cytoplasmic tau has been shown to play a role in proteasomal and mitochondrial dysfunction and aggregation, whereas nuclear tau is important in nuclear transport deficits and the loss of heterochromalin. We assessed the localization of different Tau protein variants in HT22 cells via confocal microscopy after 48 h of transfection. We found that TauS214E/T231E/S356E (Tau^3E^), TauS214A, TauT231A, TauT231E, TauS356A, and TauS356E were localized mainly in the cytoplasm, with the neuclus/cytoplasm ratio significantly lower than that of hTau (p*<*0.0001). Tau214E, TauS214A/T231A/S356A (Tau3A), and hTau are located slightly higher in the nucleus than in the cytoplasm (**Fig 1K**). A significantly greater neuclus/cytoplasm ratio was observed for Tau214E than for Tau214A (p*<*0.0001) and for Tau^3A^ than for Tau^3E^ in HT22 cells (p*<*0.0001).

**Fig. 1.**
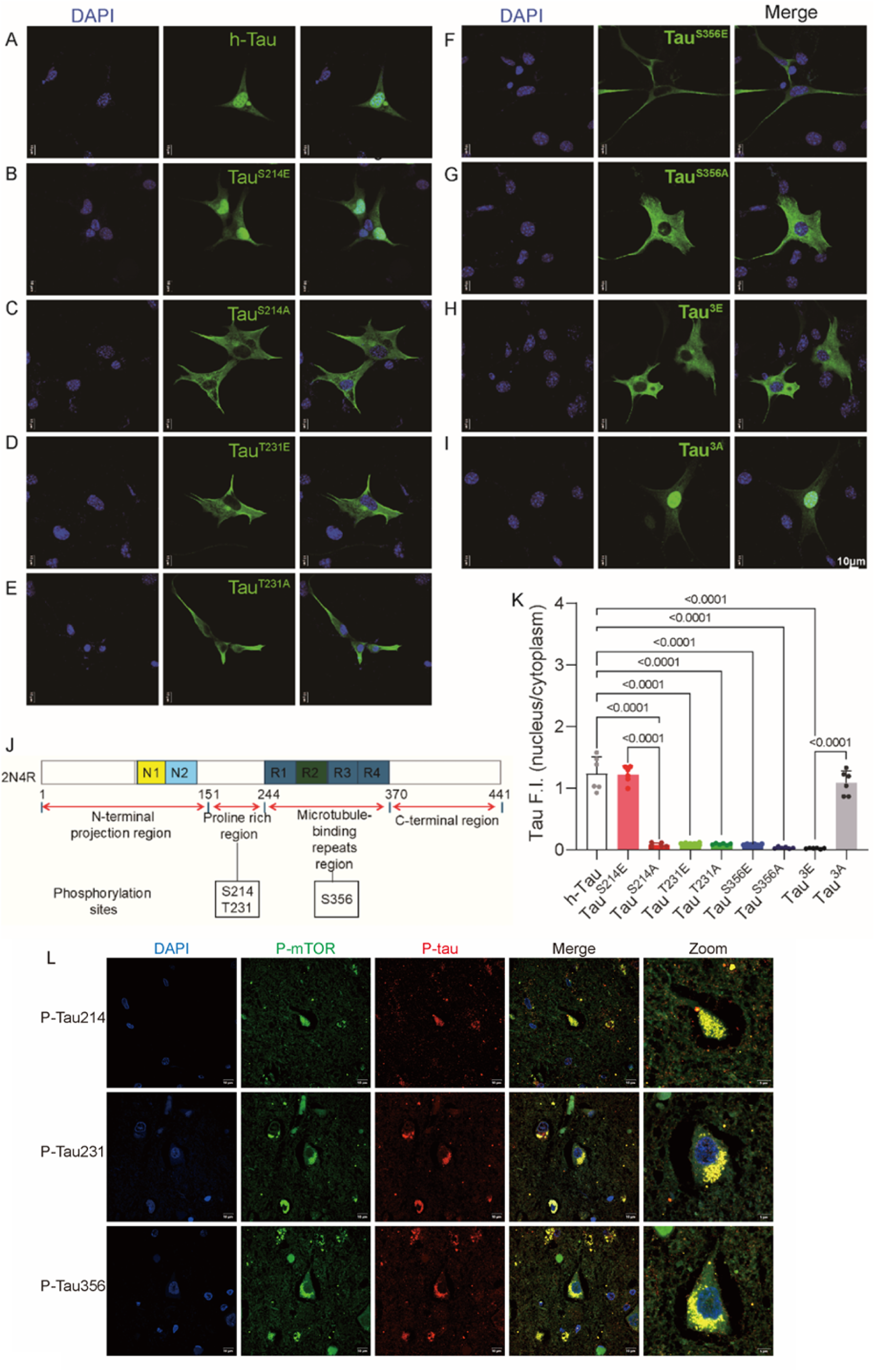
p-Tau location in the hippocampus of AD patients and HT22 cells. A-J) Altered nuclear location of tau expression in different mutants in HT22 cells. A) h-Tau; B) TauS214E; C) TauS214A; D) TauT231E; E) TauT231A; F) TauS356E; G) TauS356A; H) TauS214ET231ES356E (Tau^3E^); I) TauS214AT231AS356A (Tau^3A^); EGFP-Tau (green); nuclei counterstained with DAPI (blue). J) Location of the tau phosphorylation site. K) Different nuclear/cytoplasmic locations of tau expression in different mutants. L) Colocalization of p-mTORS2448 with p-Tau214, p-Tau231, and p-Tau356 (n=3 each). p-mTORS2448 (green); Tau (red); nuclei counterstained with DAPI (blue).

The colocalization of p-TauS214, p-T231 and p-TauS356 with p-mTORS2448 in the CA3 region of the hippocampal tissues of AD patients was observed. p-TauS214 and p-TauS356 were colocalized with mTORS2448 in the cytoplasm as well as the nucleus, and p-TauT231 was expressed mainly in the cytoplasm of the neuron stained in AD hippocampal slices (**Fig 1L**).

We chose Tau^3E^, which is simultaneously phosphorylated at these three sites and located in both the cytoplasm and neuclus, for follow-up study. Western blotting revealed that the expression levels of p-S214, p-T231 and p-S356 were similar in the control and HT22 cells transfected with h-Tau (**Fig 2A-D**). In comparison, the expression levels of p-S214 (*P =* 0.0025), p-T231 (*P <* 0.0001) and p-S356 (*P <* 0.0001) were greater in the HT22 cells overexpressing Tau^3E^ than in the control cells (**Fig 2A-D**).

**Fig. 2.**
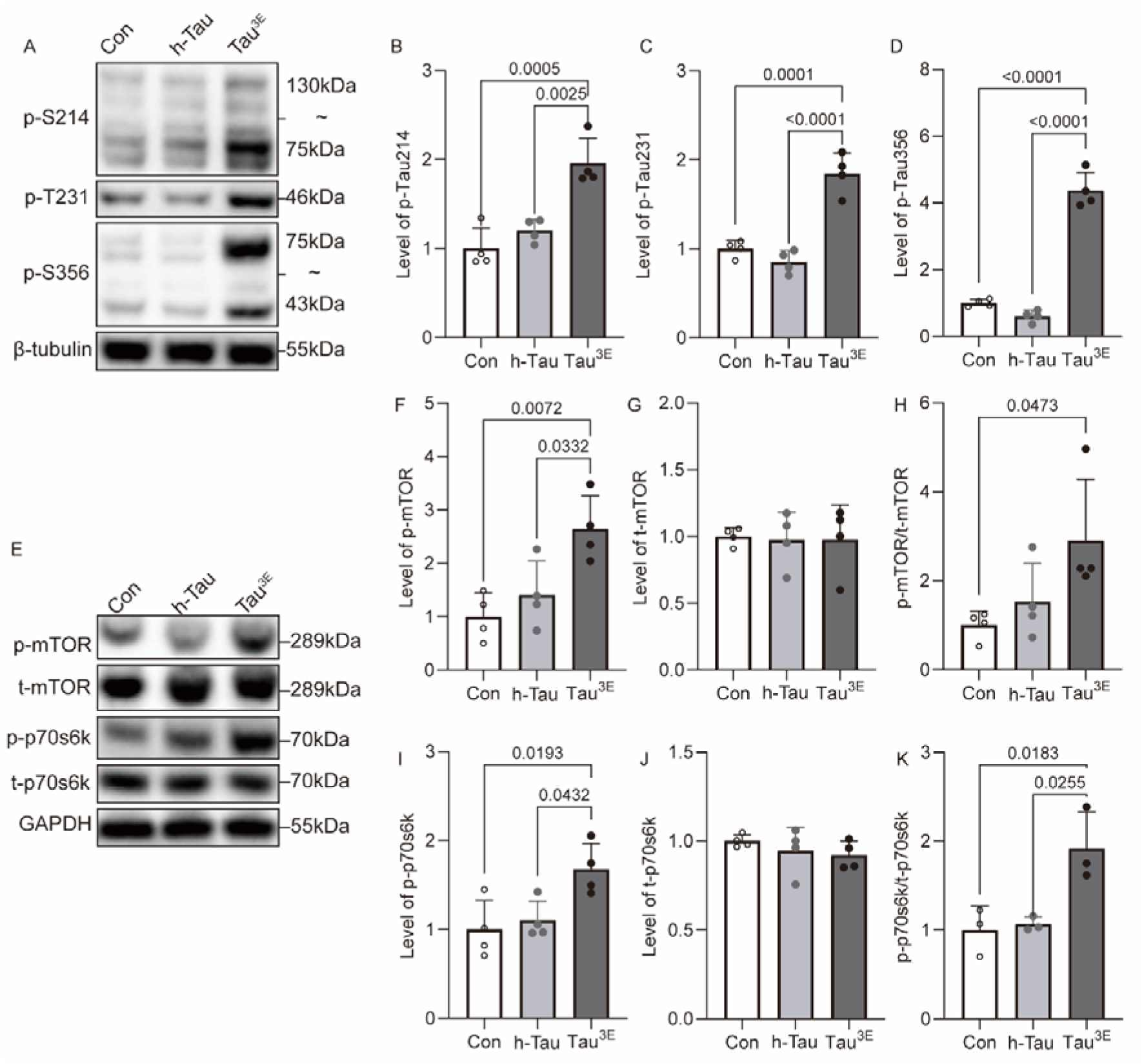
Increased tau phosphorylation and mTOR pathway-related protein expression in HT22 cells transfected with Tau^3E^. A-D) Western blot analysis of the tau proteins p-TauSer214 (B), p-TauThr231 (C), and p-TauSer356 (D). E-K): Western blot analysis of mTOR pathway-related proteins. p-mTOR protein (F), t-mTOR protein (G), p-mTOR/t-mTOR protein (H). p-p70s6k (I), t-p70s6k (J), p-p70s6k/t-p70s6k protein (K). Phosphorylated, p; total, t

### 3.2 Tau^3E^ increased the expression of mTOR pathway proteins and ROS in HT22 cells

We first assessed whether the hyperphosphorylation of Tau activated the mTOR/p70s6k pathway and induced oxidative stress. HT22 cells 48 hours after transfection with p-tau. The levels of t-mTOR, t-p70s6k, p-mTOR, p-p70s6k, mTOR/t-mTOR and p-p70s6k/t-p70s6k were comparable between the control and h-Tau groups (**Fig 2E–K**). Increased p-mTOR (*P =* 0.0332) and p-p70s6k and p-p70s6k/t-p70s6k ratios (*P =* 0.0225) were detected in HT22 cells overexpressing Tau^3E^ compared with those overexpressing h-Tau. An increased p-mTOR/t-mTOR ratio (*P =* 0.0473) was detected in the HT22 cells overexpressing Tau^3E^ compared with the control cells (**Fig 2E-K**).

To study the potential role of p-tau in oxidative stress, we used lipid peroxidation products to detect the potential of p-tau in HT22 cells after transient recombination of the tau plasmid for 48 h. 4-HNE is an electrophilic lipid peroxidation product that has various cytotoxic effects on lipid peroxidation, such as depletion of glutathione and a reduction in enzyme activity. Antibodies against 4-HNE, the DNA oxidation product 8-OHdG, and the oxidative stress-induced ROS marker NOX4 were used (**Fig. 3A-F**). Compared with h-Tau, p-Tau^3E^ increased the levels of the oxidative stress products 4-HNE (*P <* 0.0001), 8-OHdG (*P <* 0.0001) and NOX4 (*P<* 0.0001) (**Fig. 3A-F**). These results further indicate that p-Tau^3E^ can increase oxidative stress in cells. To further elucidate the mechanism underlying the effects of Tau^3E^ on oxidative stress in HT22 cells, we measured the changes in intracellular ROS levels after transient transfection with Tau. The abovementioned HT22 cells transiently transfected with tau were incubated with 2’,7’-dichlorofluorescein diacetate for 20 min. After the cells were washed with buffer, the fluorescence of DHE was detected via flow cytometry to assess the degree of tau transfection. Compared with that in the control group (h-Tau), the production level of ROS in HT22 cells after transfection with Tau^3E^ was 8.8% greater (*P =* 0.0002) (**Fig. 3G-H**). These results indicated that in HT22 cells, the expression of Tau^3E^ increased the production of ROS, further indicating that severe oxidative stress occurred in these cells.

**Fig. 3.**
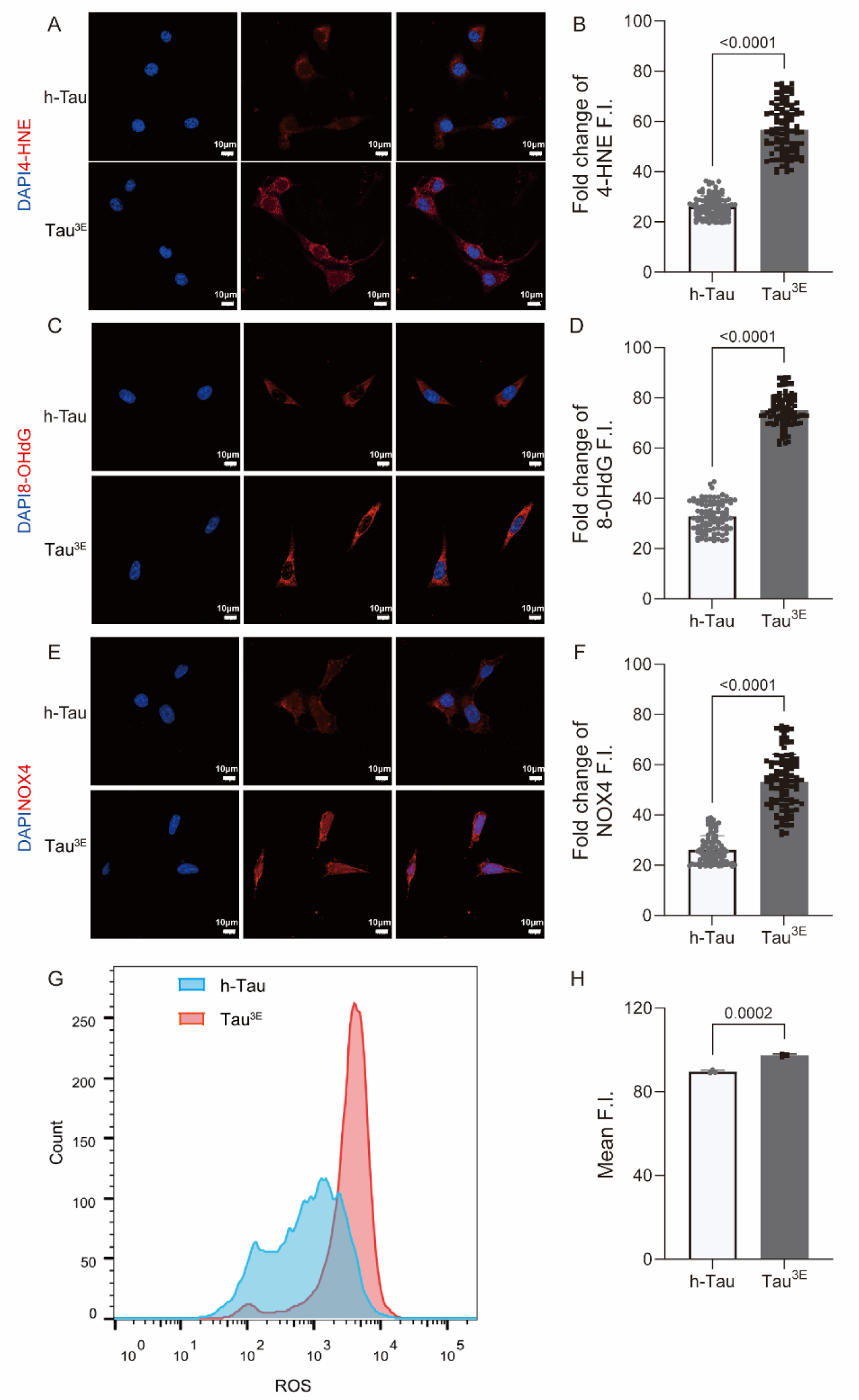
Immunofluorescence staining analysis of HT22 cells transfected with Tau^3E^. A, C, E) 4-HNE, 8-OHdG (red), and NOX4 (red) positive immunofluorescence staining. The nuclei were counterstained with DAPI (blue). B, D, F) Quantitative analysis of 4-HNE and 8-OHdG NOX4-positive fluorescence staining intensity (n=100). G, H) Representative flow cytometry image of ROS in HT22 cells overexpressing Tau^3E^ and quantitative analysis of ROS levels (n=3).

### 3.3 Tau^3E^ impaired mitochondrial structure and dynamics in HT22 cells

Next, we studied the effects of tau phosphorylation on mitochondrial homeostasis and the underlying molecular mechanism (**Fig. 4A–B**). Mitochondrial morphology was analyzed after TOMM20 staining under a confocal microscope. When more than 70% of the mitochondria in a cell are fragmented, the cell can be identified as having mitochondrial fragmentation ^[48]^. Among the HT22 cells transfected with Tau^3E^, the percentage of cells with fragmented mitochondria increased from 17.3% in the h-Tau group to 69.4% (*P =* 0.0001). These results indicated that in the HT22 cells, compared with that in the h-Tau group, the expression of Tau^3E^ severely damaged mitochondrial structure (**Fig 4A-B**).

**Fig. 4.**
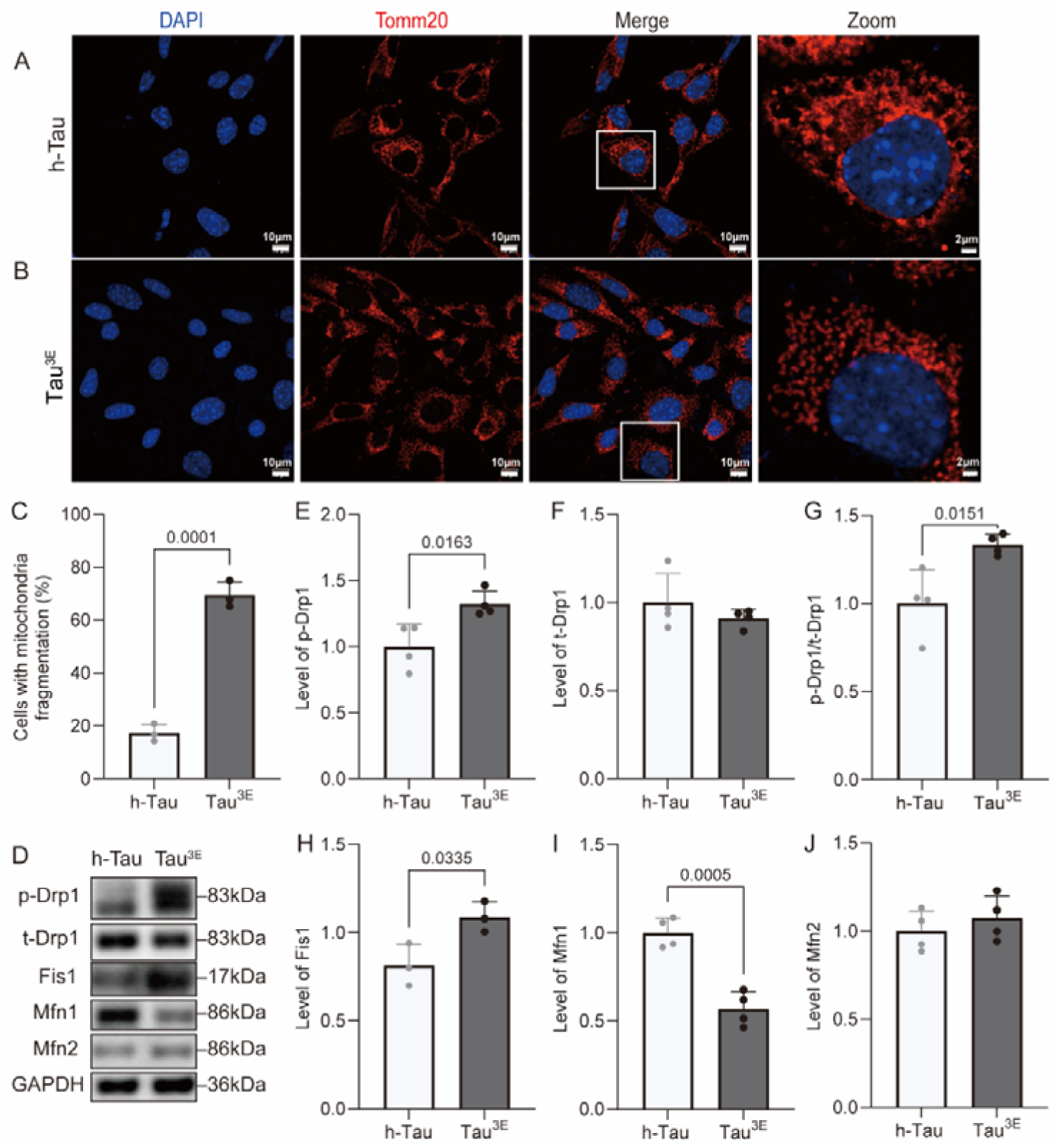
Increased mitochondrial damage, mitochondrial division and fusion-related protein expression in HT22 cells overexpressing Tau^3E^. A-C): Representative immunofluorescence images of TOMM20 (red) in HT22 cells overexpressing Tau^3E^ and h-Tau and greater mitochondrial fragmentation in Tau^3E-^ and h-Tau-overexpressing cells (C). Nuclei counterstained with DAPI (blue). D-J): Western blot analysis of mitochondrial division- and fusion-related proteins in HT22 cells overexpressing Tau^3E^ and h-Tau; p-Drp1 (E), t-Drp1 (F), p-Drp1/t-Drp1 (G), Fis1 (H), Mfn1 (I), and Mfn2 (J).

We assessed the levels of the mitochondrial fission fusion proteins t-Drp1, p-Drp1, Fis1, Mfn1, and Mfn2 in HT22 cells overexpressing Tau^3E^ via Western blotting at 48 h. No difference in the levels of the mitochondrial fission-fusion-related proteins t-Drp1 and Mfn2 was detected between the p-Tau^3E^ group and the h-Tau group (**Fig 4C, F, I**). Compared with those in the h-Tau group, increased levels of the mitochondrial fission-related proteins p-Drp1 (*P =* 0.0076), p-Drp1/t-Drp1 (*P =* 0.0151) and Fis1 (*P =* 0.0335) were detected in the p-Tau^3E^ group (**Fig 4C, D, E, G**). A lower level of the mitochondrial fusion-related protein Mfn1 (*P =* 0.0005) was detected in the p-Tau^3E^ group than in the h-Tau group (**Fig 4C, H**).

### 3.4 Rapamycin alleviated impaired cognitive behavior and neuronal loss in Tau^3E^ mice

To generate a mouse model overexpressing Tau^3E^ in the CA3 region of the hippocampus, AAV9 adeno-associated viral vectors were ligated with Tau^3E^ plasmids to construct recombinant adeno-associated virus expression vectors in vitro. AAV-Tau^3E^ with green fluorescent protein and AAV-control were injected into the CA3 region of the hippocampus of 2-month-old C57BL/6 male mice via the stereotaxic injection technique. The location of green fluorescence in frozen sections of mouse brain tissue confirmed the injection of the viral vector into the CA3 area of the mouse hippocampus (**Fig. 5A**).

**Fig. 5.**
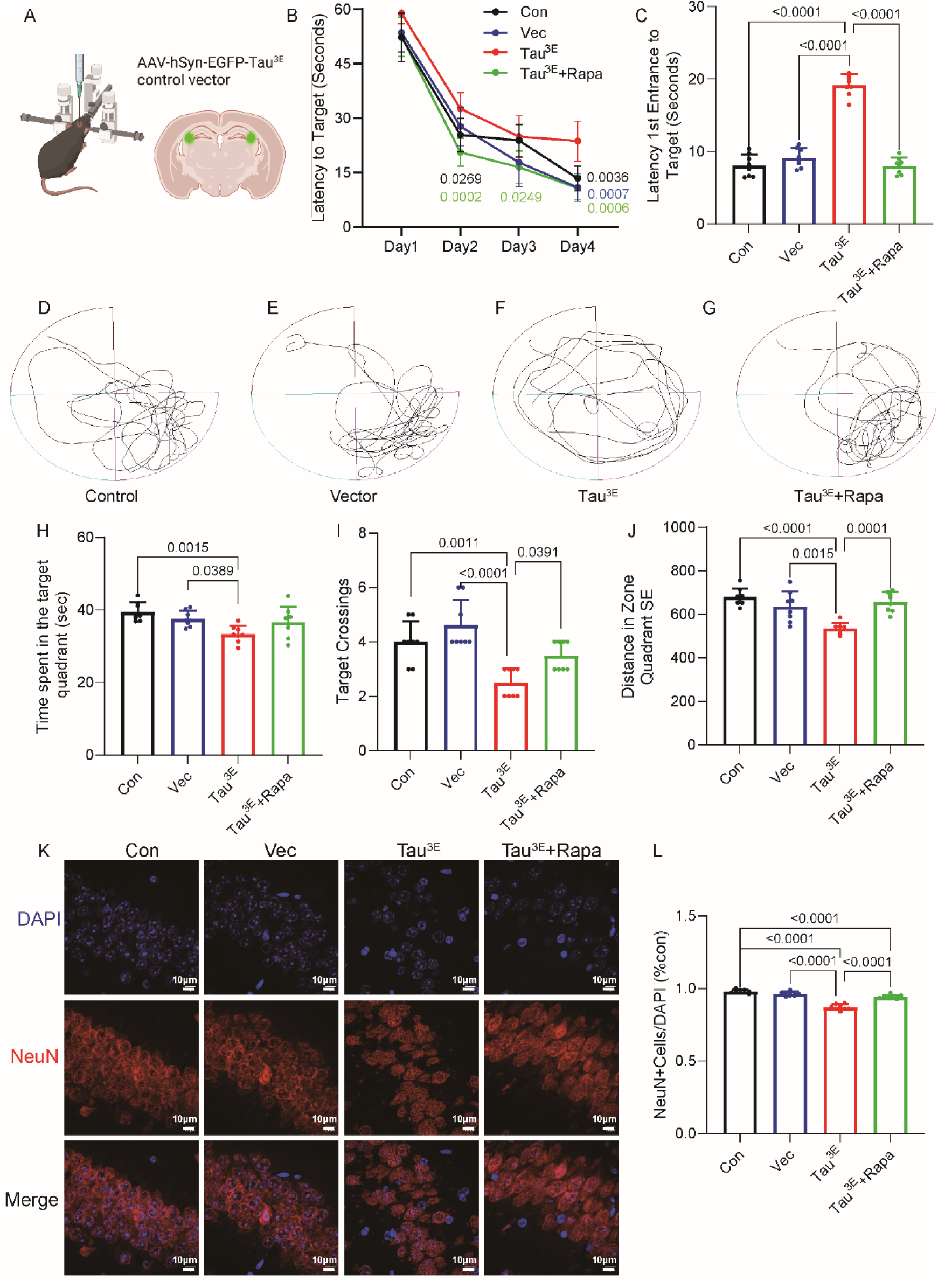
Rapamycin (i.p.) treatment ameliorated cognitive impairment and reduced the number of neurons in Tau^3E^ mice. A) Schematic representation of the bilateral injection of AAV9 adenovirus overexpressing Tau^3E^ or the control construct in the CA3 region of the hippocampus in C57BL/6 mice. B): Escape latency of the mice. C) Time taken by the mice to first reach the platform; DLJG) track of swimming; H) time spent by the mice in the target quadrant; I) number of times the mice crossed the platform. J: Distance traveled by the mice in the target quadrant (n=8). K) NeuN (red) immunofluorescence staining images of the CA3 area in the hippocampus nuclei counterstained with DAPI (blue). L) Quantification of NeuN-positive cells/cell nuclei (DAPI) (n=3).

To test whether rapamycin can rescue cognitive dysfunction in the four groups (control, vector, Tau^3E^ and Tau^3E^ + rapa), we intraperitoneally injected 1.5 mg/kg rapamycin three times a week into the Tau^3E^ + rapa group. The Morris water maze test was performed after 1 week. The learning outcomes of the mice were assessed on day 5 after 4 days of learning. In the escape latency test, there was no difference in the performance of the four groups of mice on the first day. On the second day, the latency of the mice to reach the platform was greater in the Tau^3E^ group than in the control group (*P =* 0.0169) and the Tau^3E^ + rapa group (*P<* 0.0001). On the third day, the latency of the mice to reach the platform was greater in the Tau^3E^ group than in the Tau^3E^ + rapa group (*P=* 0.0196); on the fourth day, the latency of the mice to reach the platform was greater in the Tau^3E^ group than in the control group (*P=* 0.0011), the vector group (*P<* 0.0001) and the Tau^3E^ + rapa group (*P<* 0.0001) (**Fig. 5B–G**). The time spent in the target quadrant was lower in the Tau^3E^ group than in the control group (*P =* 0.0015) and the vector group (*P =* 0.0389) (**Fig. 5H**). The number of platform crossings was lower in the Tau^3E^ group than in the control group (*P =* 0.0011), vector group (*P <* 0.0001), and Tau^3E^ + rapa group (*P =* 0.0391) (**Fig. 5I**). The distance in the zone was shorter in the Tau^3E^ group than in the control group (*P<* 0.0001), vector group (*P=*0.0015), and Tau^3E^ + rapa group (*P =* 0.0001) (**Fig. 5J**). These results suggest that the overexpression of Tau^3E^ impaired cognitive ability in mice and that rapamycin (i.p.) treatment reversed these cognitive impairments.

Paraffin sections of mouse brain tissue were labeled with a NeuN antibody, and changes in the fluorescence intensity in the CA3 region of the hippocampus were observed via multiplex immunofluorescence staining. Compared with that in the control group, the number of neuronal cells in the Tau^3E^ group was 11% lower (*P* <0.0001), which is consistent with earlier results(*32*). The number of neuronal cells was 8% lower in the Tau^3E^ group than in the Tau^3E^ + rapa group (*P* <0.0001) (**Fig. 5K, L**).

### 3.5 Rapamycin reduced p-Tau expression in Tau^3E^ mice

Next, we assessed whether there was a difference in the levels of p-tau proteins after treatment with rapamycin. The protein expression levels of p-S214, p-T231, and p-S356 in the control group and vector group were not significantly different in the hippocampus or in the cortex. Compared with the control and vector groups, the Tau^3E^ group presented increased p-S214 (*P<* 0.0001, *P<* 0.0001), p-T231 (*P<* 0.0001, *P<* 0.0001) and p-S356 (*P<* 0.0001*, P<* 0.0001) levels in the hippocampus, as well as in the cortex in p-S214 (*P=*0.0003, *P=*0.0016), p-T231 (*P=*0.0002, *P=* 0.0117) and p-S356 (*P=*0.0004*, P=*0.0008). Rapamycin treatment significantly reduced the levels of p-S214 (*P=* 0.0003; *ns*), p-T231 (*P<* 0.0001; *P=* 0.0308), and p-S356 (*P=*0.0005*; P=* 0.0066) in the hippocampus and cortex (Tau^3E^ group vs Tau^3E^+rapa, **Fig. 6A–H**). The expression of EGFP-Tau^3E^ was further validated by confocal microscopy of coronal brain slices from mouse brains, which revealed fluorescence in the hippocampus (particularly CA3) as well as in the cortical region (**Fig. 6I**).

**Fig. 6.**
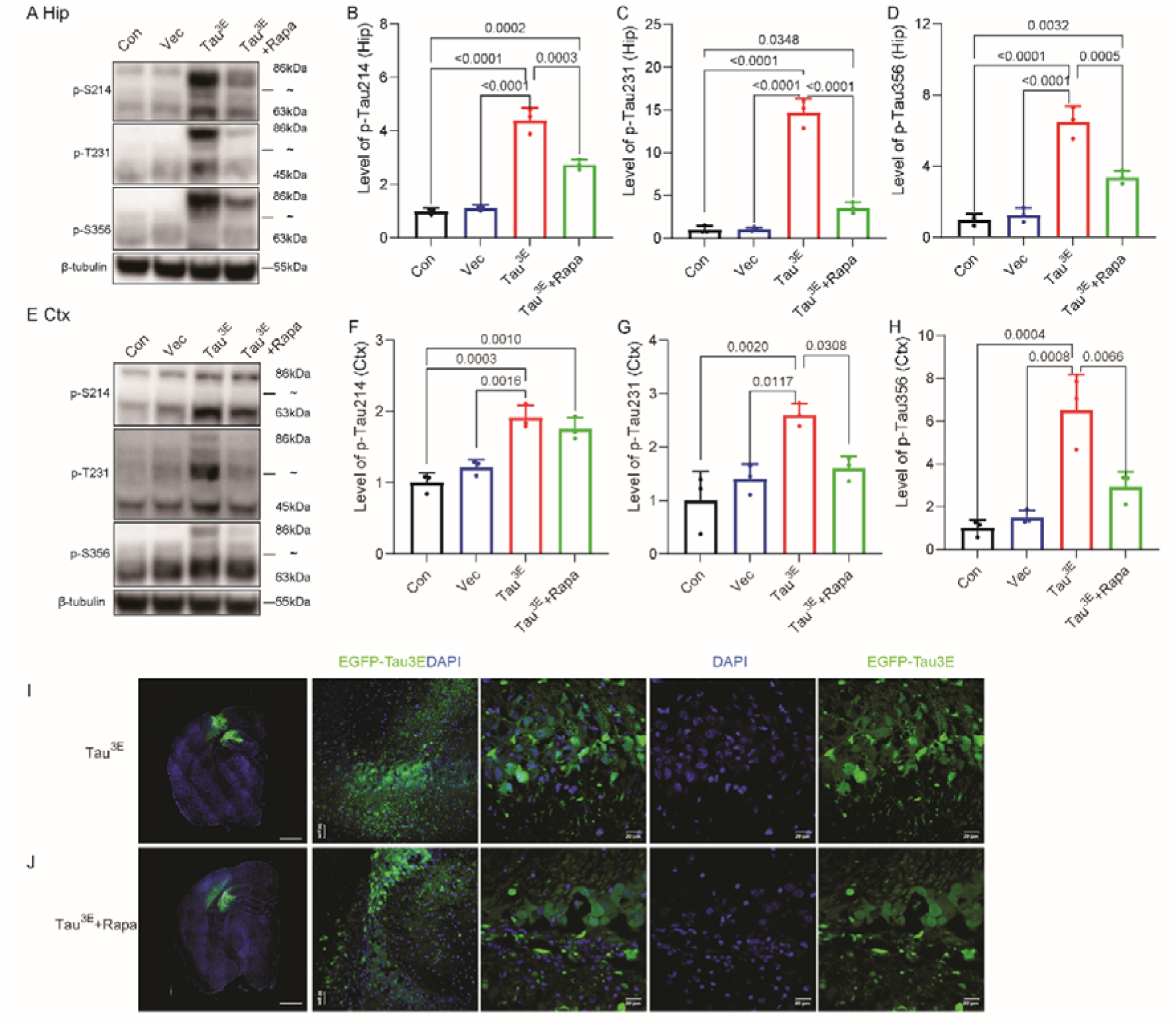
Rapamycin treatment (i.p.) reversed the increase in Tau expression in the brains of Tau^3E^ mice. A–H) Western blot analysis of Tau-related proteins in the hippocampus and cortex; Hippocampal and cortical levels of B, F) p-TauSer214; C, G) p-TauThr231; D, H) p-TauSer356 (n=3). I, J) Expression of EGFP-Tau^3E^ (green) in the mouse brain after injection of AAV9 hSyn-EGFP-Tau^3E^ in the CA3 region of Tau^3E^ mice. The nuclei were counterstained with DAPI (blue).

### 3.6 Rapamycin restores metabolic profile, signaling microenvironment and neurotransmitter homeostasis in Tau^3E^ mice

To evaluate the effect of rapamycin on tau-induced metabolic dysfunction, we assessed the metabolomic profiles of the brain tissue from the Tau^3E^ and Tau^3E^+rapa groups. Distinct cluster separation was observed between the two groups by PCA, suggesting that rapamycin treatment significantly altered the neurochemical environment of the hippocampi of Tau^3E^ mice (**Fig. 7A**). As shown in the volcano map, we further pinpoint the specific metabolites modulated by rapamycin in Tau^3E^ mice, including a total of 3732 differentially abundant metabolites, of which 716 increased and 360 decreased (**Fig. 7B**). KEGG topology analysis revealed that multiple metabolic pathways were enriched in the Tau^3E^+rapa group compared with the Tau^3E^ group, particularly glycerophospholipid metabolism, unsaturated fatty acid biosynthesis, purine metabolism, and choline metabolism in cancer and insulin resistance (**Fig. 7C**).

**Fig. 7.**
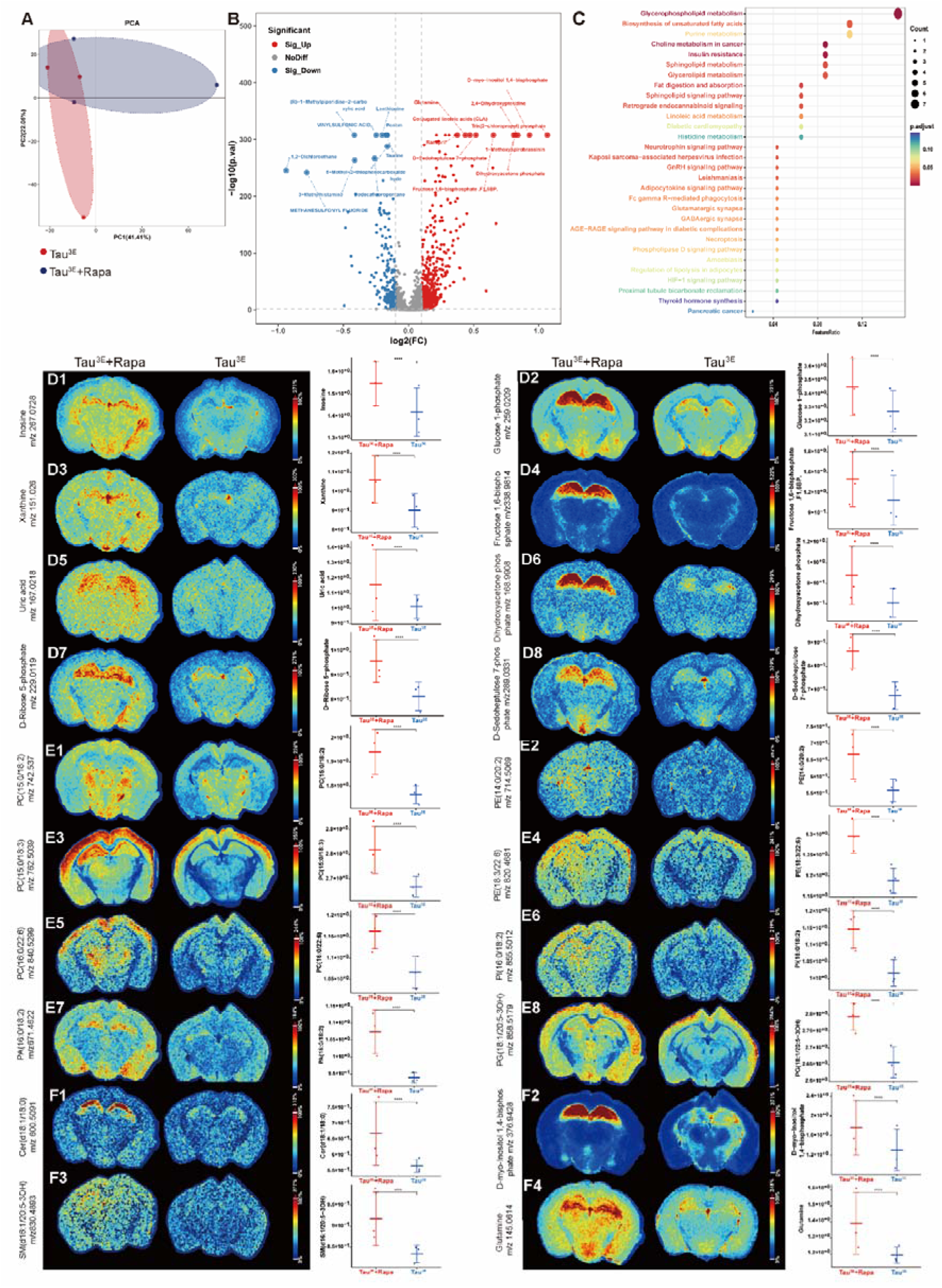
Identification of in situ small-molecule metabolites involved in the protective effect of rapamycin by AFADESI-MSI in the brains of Tau^3E^ mice. A) Principal component analysis (PCA) of metabolites in the hippocampi of the Tau^3E^ and Tau^3E^+rapa group. n=3 per group. B) Volcanic map showing different metabolites in the hippocampi of the Tau^3E^ and Tau^3E^+rapa groups. C) KEGG enrichment analysis of different metabolites in the hippocampi of the Tau^3E^ and rapamycin-treated groups. D) Different metabolites of the inosine-ppp-glycolysis axis, including inosine (m/z 267.0728) (D1), glucose 1-phosphate (m/z 259.0209) (D2), xanthine (m/z 151.026) (D3), fructose 1,6-bisphosphate (m/z 338.9814) (D4), uric acid (m/z 167.0218) (D5), dihydroxyacetone phosphate (m/z 168.9908) (D6), D-ribose 5-phosphate (m/z 229.0119) (D7) and D-sedoheptulose 7-phosphate (m/z 289.0331) (D8) in the hippocampi of the Tau^3E^ and Tau^3E^+rapa groups. E) Different metabolites of the lipidome axis including phosphatidylcholines (PCs, E1, E3, E5), 15:0/18:2 (m/z 742.537), 15:0/18:3 (m/z 762.5039), and 16:0/22:6 (m/z 840.5299); phosphatidylethanolamines (PEs, E2, E4), 14:0/20:2 (m/z 714.5069), and 18:3/22:6 (m/z 820.4681); phosphatidylinositols (PIs, E6), 16:0/18:2 (m/z 855.5012); phosphoric acid (PA, E4) 16:0/18:2 (m/z 671.4622); phosphatidylglycerol (PG, E8) 18:1/20:5 (m/z 858.5179), in the hippocampi of the Tau^3E^ and Tau^3E^+rapa groups. F) Different metabolites involved in the signaling microenvironment and neurotransmitter homeostasis including ceramide (Cer) (d18:1/18:0) (m/z 600.5091)(F1), D-myo-inositol 1,4-bisphosphate (m/z 376.9428)(F2), sphingomyelin (SM) d16:1/20:5 (m/z 830.4893)(F3), and glutamine (m/z 145.0614)(F4) in the hippocampi of the Tau^3E^ and Tau^3E^+rapa groups.

Using air-flow-assisted desorption electrospray ionization (AFADESI)-MSI analysis, we integrated spatial and quantitative data to characterize metabolic profiles in the hippocampus of mice. Rapamycin treatment significantly reversed this metabolic decay by increasing the in situ concentration of key catabolites in the inosine-pentose phosphate pathway (PPP)-glycolysis metabolic axis. Specifically, rapamycin promoted robust accumulation of purine catabolites, including inosine (m/z 267.0728), xanthine (m/z 151.026), glucose 1-phosphate (m/z 259.0209) and uric acid (m/z 167.0218). This upregulation was closely coordinated with the activation of the PPP, as evidenced by the enriched spatial distribution of D-ribose 5-phosphate (m/z 229.0119) and D-sedoheptulose 7-phosphate (m/z 289.0331) in the hippocampus (**Fig. 7D**). Furthermore, the compensatory flux through the PPP appeared to improve energy homeostasis, as evidenced by the significant restoration of glycolytic markers such as fructose 1,6-bisphosphate (m/z 338.9814) and dihydroxyacetone phosphate (m/z 168.9908) (**Fig. 7D**).

In addition, rapamycin treatment reversed the decrease in the level of the structural lipidome axis in the hippocampus (Tau^3E^+rapa vs. Tau^3E^ mice) by restoring a broad spectrum of essential phospholipids. Specifically, we observed in situ recovery of phosphatidylcholines (PCs), including 15:0/18:2 (m/z 742.537), 15:0/18:3 (m/z 762.5039), and 16:0/22:6 (m/z 840.5299), which are critical for maintaining membrane fluidity (**Fig. 7E**). This restoration was closely associated with the upregulation of phosphatidylethanolamines (PEs), e.g., 14:0/20:2 (m/z 714.5069), and 18:3/22:6 (m/z 820.4681); and phosphatidylinositols (PIs), e.g., (16:0/18:2) (m/z 855.5012), which are essential for synaptic signaling and cellular metabolism. Furthermore, rapamycin treatment increased the levels of signaling and precursor lipids, including phosphoric acid (PA) (16:0/18:2) (m/z 671.4622) and phosphatidylglycerol (PG) (18:1/20:5) (m/z 858.5179), in the mouse hippocampus (**Fig. 7E**). We further integrated spatial profiles of sphingolipid signaling, inositol-related second messengers, and amino acid metabolites. Rapamycin treatment counteracted this deterioration through systematic reprogramming of the hippocampal signaling microenvironment, including sphingolipid mediators, such as ceramide (Cer) (d18:1/18:0) (m/z 600.5091) and sphingomyelin (SM) (d16:1/20:5) (m/z 830.4893) (**Fig. 7F**). This was spatially associated with the upregulation of the inositol signaling pathway, e.g., D-myo-inositol 1,4-bisphosphate (m/z 376.9428), restoring the capacity for intracellular communication. Furthermore, rapamycin increased the local abundance of glutamine (m/z 145.0614), a critical component for maintaining the glutamate cycle in the central nervous system (**Fig. 7F**).

### 3.7 Rapamycin reduced mTOR pathway protein expression in Tau^3E^ mice

Next, we quantified whether mTOR pathway activity is affected in tau mice and in response to rapamycin treatment in the cortex and hippocampus. Compared with the Tau^3E^ group, the Rapa treatment (i.p.) reversed the Tau^3E^-induced increase in the p-mTOR (*P=*0.0296, *P=*0.0260) and p-mTOR/t-mTOR (*P=*0.0112, *P=*0.0080) levels in the hippocampi and cortices of the mice in the Tau^3E^ +Rapa group (**Fig. 8C, E, I, K**). Western blot analysis revealed that the protein expression levels of t-mTOR and t-p70s6k did not significantly differ among the four groups of mice (**Fig. 8E, G, J, M**). Compared with the control and vector groups, the Tau^3E^ group presented higher p-mTOR (*P<* 0.0008, *P=*0.0001) and p-mTOR/t-mTOR (*P<* 0.0001, *P=0.0004*) levels in the hippocampus and p-mTOR (*P=* 0.0002, *P=*0.0002) and p-mTOR/t-mTOR (*P<* 0.0001, *P<* 0.0001) levels in the cortex. Compared with the control and vector groups, the Tau^3E^ group presented higher levels of p-p70s6k (*P=* 0.0038, *P=*0.0057) in the hippocampus and (*P=* 0.0023, *P=*0.0064) in the cortex and p-p70s6k/t-p70s6k (*ns*, *P=* 0.0114) in the cortex. Compared with the Tau^3E^ group, Rapa treatment (i.p.) reversed the Tau^3E^-induced increase in the levels of p-p70s6k (*P=*0.0250, *P=*0.0132) and p-p70s6k/t-p70s6k (*ns*, *P=*0.0395) in the hippocampi and cortices of mice in the Tau^3E^ +Rapa group (**Fig. 8F–H, L-N**).

**Fig. 8.**
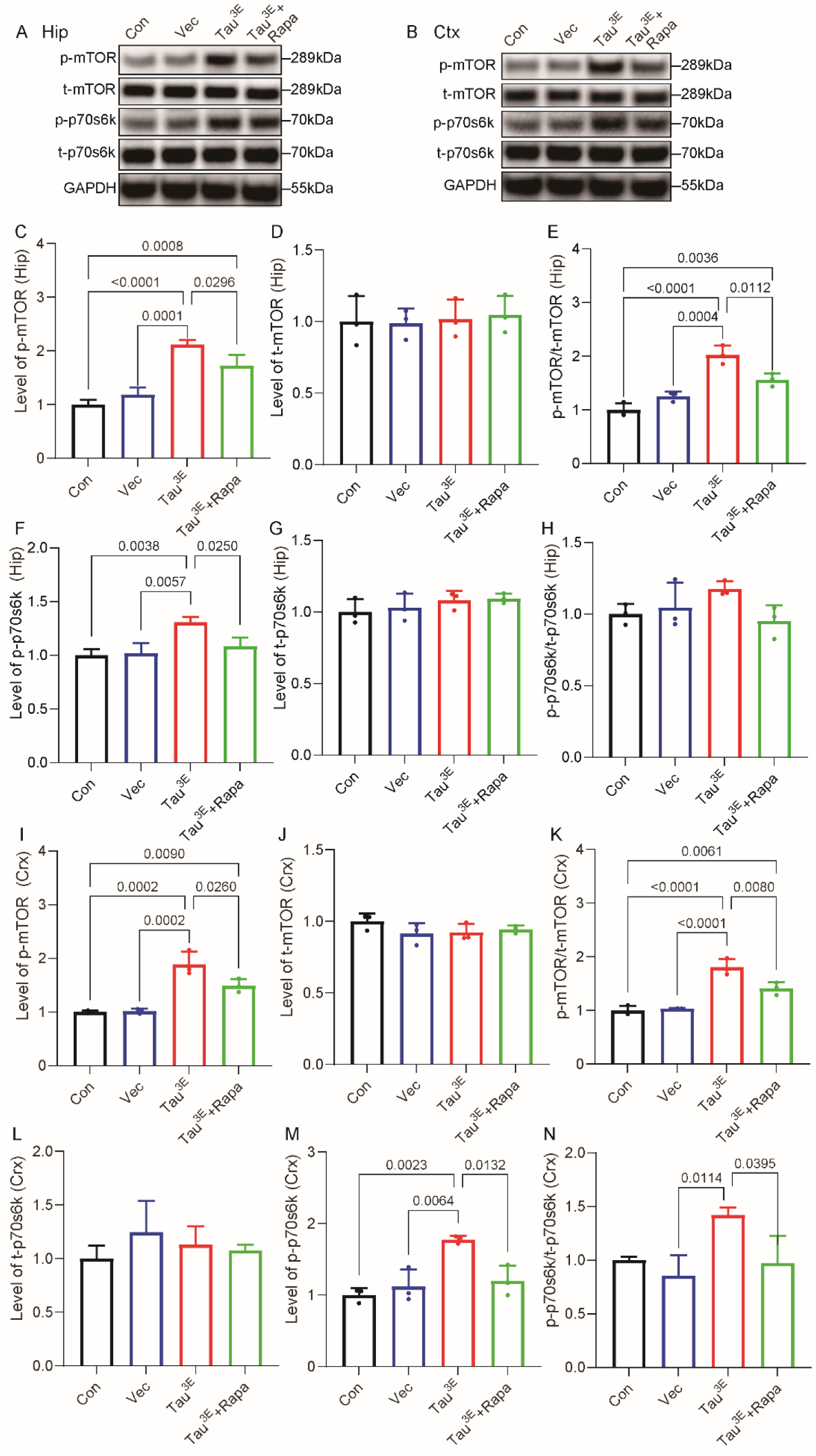
Rapamycin treatment (i.p.) reversed the increase in the levels of mTOR pathway-related proteins in the brains of Tau^3E^ mice. A, B) Western blot analysis of the levels of mTOR pathway-related proteins in the hippocampus and cortex of Tau^3E^ and Tau^3E^+rapa groups. Hippocampal and cortical levels of C, I) p-mTOR; D, J) t-mTOR; E, K) p-mTOR/t-mTOR ratio; F, L) p-p70s6k protein; G, M) t-p70s6k; H, N) p-p70s6k/t-p70s6k ratio (n=3).

### 3.8 Rapamycin reduced the increase in the oxidative stress products 4-HNE, 8-OHdG and NOX4 in Tau^3E^ mice

To investigate the alterations in oxidation parameters caused by Tau^3E^ and rapa treatment, paraffin sections of mouse brain tissue were labeled with 4-HNE, 8-OHdG and NOX4 antibodies. There was no significant difference in the fluorescence intensity of 4-HNE, 8-OHdG, or NOX4 between the control group and the vector group. Compared with those in the control group and the vector group, the intensities of positive staining for the oxidation indicators 4-HNE (*P* <0.0001, *P* <0.0001), 8-OHdG (*P* <0.0001, *P* <0.0001), and NOX4 (*P* <0.0001, *P* <0.0001) increased in the hippocampus (CA3) of mice in the Tau^3E^ group. Compared with the Tau^3E^, the Rapa treatment reduced the Tau^3E-^induced increase in the intensities of 4-HNE (*P* <0.0001), 8-OHdG (*P* <0.0001), and NOX4 (*P* <0.0001) in the hippocampi of the Tau^3E^ +rapa group (**Fig 9A–F**).

**Fig. 9.**
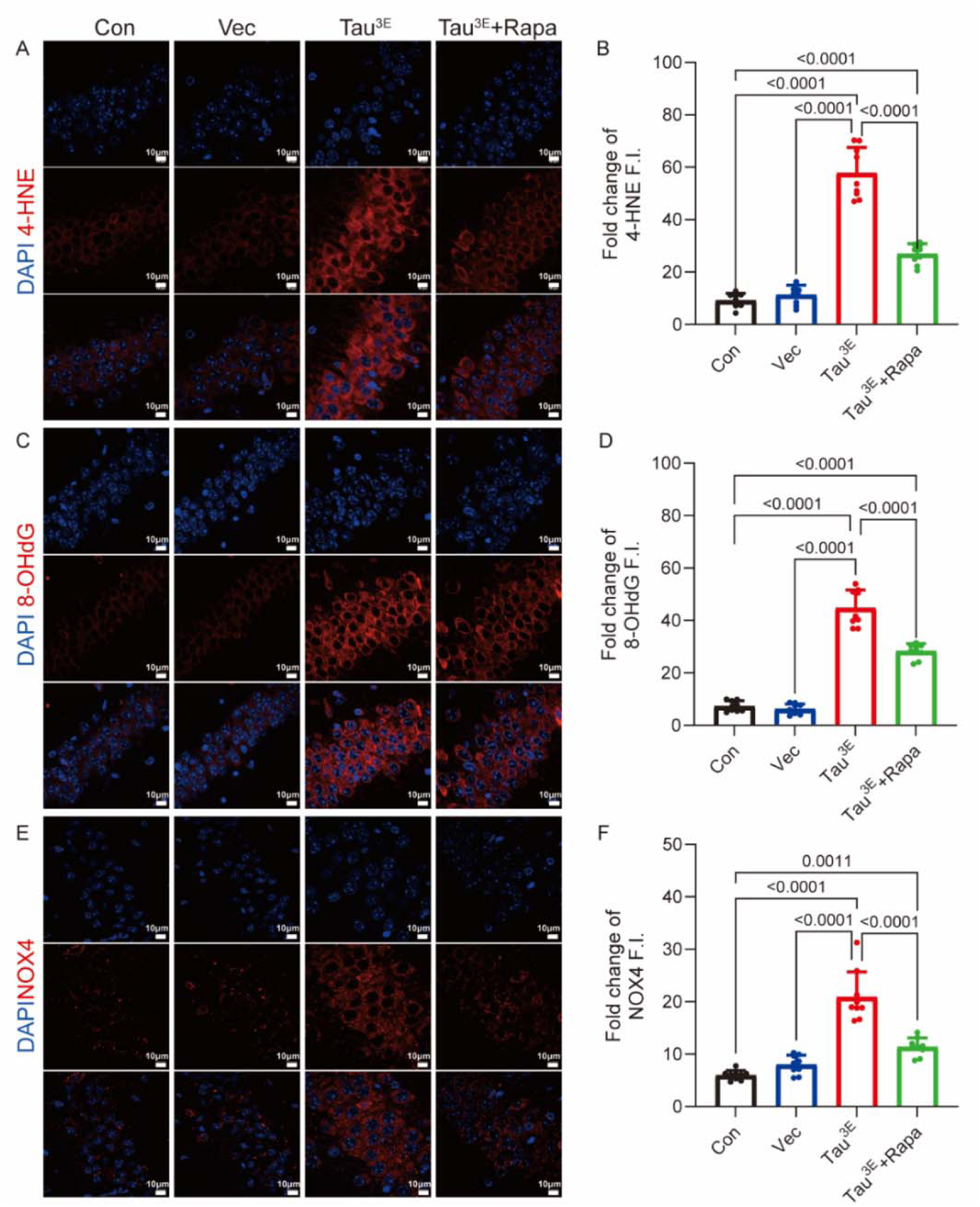
Rapamycin treatment (i.p.) reversed the increase in the oxidation indices 4-HNE, 8-OHdG and NOX4 in the brains of Tau^3E^ mice. Immunofluorescence images and quantification of 4-HNE (A, B), 8-OHdG (C, D), and NOX4 levels (E, F). 4-HNE (red), 8-OHdG (red), and NOX4 (red) in the cortex and hippocampi of the Tau^3E^ and Tau^3E^+rapa groups; nuclei were counterstained with DAPI (blue).

### 3.9 Rapamycin improved impaired mitochondrial dynamics in Tau^3E^ mice

Proteins were extracted from mouse cortical and hippocampal tissues, and the levels of the mitochondrial fission-fusion proteins t-Drp1 and p-Drp1 and the mitochondrial fusion-related proteins Mfn1 and Mfn2 were detected via western blotting. The t-Drp1 protein expression levels in the hippocampus and cortex of mice in the control, vector, Tau^3E^, and Tau^3E^ + rapa groups were comparable. Compared with the control group, the Tau^3E^ group presented increased expression of p-Drp1/t-Drp1 and p-Drp1 in the hippocampus (*P* <0.0001, *P* <0.0001) and in the cortex (*P* <0.0003, *P* <0.0001; **Fig. 10A-D, G-J**). Compared with the vector group, the Tau^3E^ group presented increased expression of p-Drp1/t-Drp1 and p-Drp1 in the hippocampus (*P* <0.0001, *P* <0.0001) and in the cortex (*P* <0.0001, *P* <0.0003; **Fig. 10A-D, G-J**). Compared with Tau^3E^, Rapa treatment (i.p.) reversed the Tau^3E^-induced increase in the p-Drp1/t-Drp1 (*P* <0.0001, *P* <0.0161) and p-Drp1 (*P* <0.0001, *P* <0.0055) ratios in the hippocampi and cortices of mice in the Tau^3E^ +Rapa group (**Fig. 10A–D, G-J**).

**Fig. 10.**
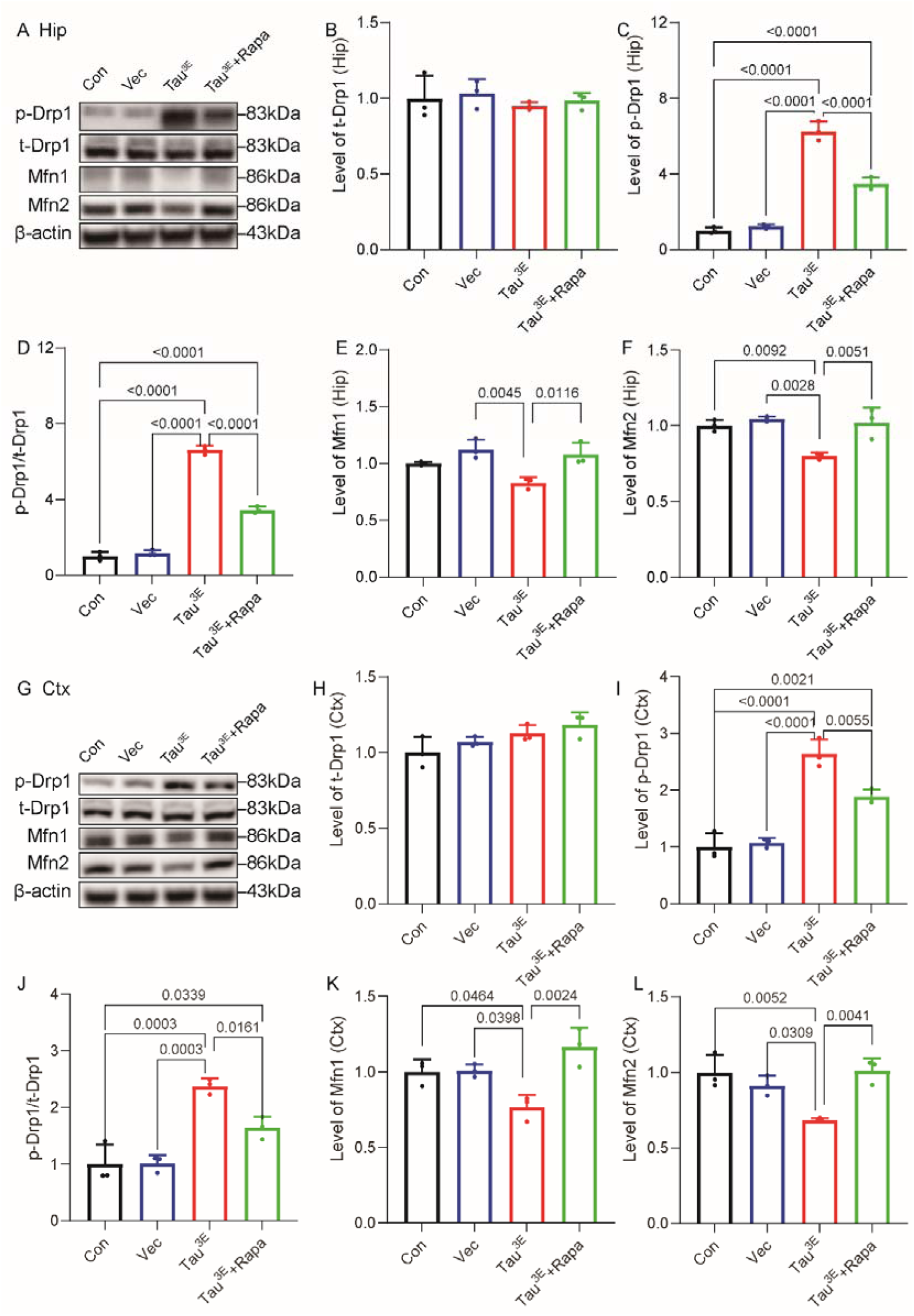
Rapamycin treatment (i.p.) reversed the increase in mitochondrial fission and fusion protein expression in the brains of Tau^3E^ mice. A–L) Western blot analysis of t-Drp1 (B, H), p-Drp1 (C, I), and the p-Drp1/t-Drp1 ratio (D, J); K) Mfn1; L) Mfn2 in the cortex and hippocampi of the Tau^3E^ and Tau^3E^+rapa groups.

In the hippocampus, the expression of Mfn1 did not significantly change in the control group but was reduced in the vector group (*P* = 0.0045) compared with that in the Tau^3E^ group, whereas the expression of Mfn2 was reduced (*P* = 0.0092; *P* = 0.0028) in the Tau^3E^ group compared with that in the control group or the vector group. In the cortex, the expression of both Mfn1 and Mfn2 was lower in the Tau3E group than in the control (*P* = 0.0464*, P* = 0.0052) and vector groups (*P* = 0.0398*, P* = 0.0309) (**Fig. 10A, E-G, K-L**). Compared with Tau^3E^, Rapa treatment (i.p.) reversed the decreases in the levels of Mfn1 (*P* = 0.0116, *P* = 0.0024) and Mfn2 (*P* = 0.0051*, P* = 0.0041) in the hippocampi and cortices of mice in the Tau^3E^ +Rapa group (**Fig. 10A, E-G, K-L**).

## 4 Discussion

Here, we demonstrated a causal relationship between tau phosphorylation at the S214, T231 and S356 sites (Tau^3E^) and mTOR upregulation. Using cellular and animal models. We revealed links between tau phosphorylation and associated oxidative stress, mitochondrial dysfunction, and cognitive impairment.

Our previous studies reported that a more than 10-fold increase in the number of AD brains is selectively associated with the p-tau epitopes at the Ser202, Thr231 and Thr231/Ser235 sites (*44*). Pseudophosphorylating tau at the proline rice region and repeat regions (Thr-212, Thr-231, and Ser(P)-262) leads to not only the loss of its normal function but also the gain of toxic activity that causes disruption of the microtubule network and cell death (*43*). Tau in SH-SY5Y cells is hyperphosphorylated at Ser214, Thr231/Ser235, Ser262, and Ser396/404 (*45*) and fully inhibits microtubule assembly in vitro, although approximately 3 times more tau in SH-SY5Y cells is required to achieve the same inhibition as AD p-tau. Our previous results from in vitro phosphorylation assays and cellular models revealed that mTOR phosphorylates tau at Ser214, Thr231, and Ser356, a similar group of epitopes shared by Akt and S6K (*46, 47*) that exclusively regulate the phosphorylation sites of tau in the proline rice region and repeat regions, resulting in inhibition of microtubule binding (*48*). These signals are critical for the conversion of tau into a toxic molecule. Tau molecular diversity contributes to clinical heterogeneity in AD. p-TauT231/S235 in AD brain lysates has been shown to increase tau seeding efficiency (*49*). In addition, the nuclear translocation of tau is a leading step in the pathological process through P53 stabilization and nucleolar dispersion(*50*). We detected differences in the nuclear and cytoplasmic localization of different tau variants, such as hTau, p-TauS214E, and p-Tau3A, in the nucleus compared with p-TauS214A, p-TauS231E, p-TauS231A, p-TauS356E, p-TauS356A, and p-Tau3E in HT22 cells. In comparison, in the human brain, p-TauS214 and p-TauS356 were detected in both the nucleus and the cytoplasm. Different tau phosphorylation sites affect the function of tau: The phosphorylation of the tau protein is regulated by the activities of protein kinases and protein phospholipases. TauS214 is an AD-specific phosphorylation site (*51*) that promotes neurotoxicity in vivo(*52*) *and* was shown to be one of the earliest significantly increased forms of neuronal phospho-tau. A recent study also revealed a protective effect of pTauSer214 on the inhibition of tau aggregation (*53*). Plasma p-tau231 and p-tau217 are state markers of Aβ pathology in preclinical AD (*54, 55*). S205, S212, S214 and S235 affect the binding of microtubules, eventually leading to neurodegeneration(*56, 57*). p-TauT231 is a promising CSF and plasma biomarker in AD (*58, 59*). p-TauT231 can decrease the affinity of tau to promote microtubule assembly and inhibit microtubule interactions (*60, 61*). p-TauT231 is present in the soluble fraction in pretangles (*62*) and accumulates at the postsynaptic density in early AD (*63–65*). Site-specific phosphorylation of tau GFP-tagged T231/S235 has been shown to impact mitochondrial function and the response to stressors (*66*). p-TauS356 is associated with AD pathology, increases with Braak stage and is present in neurofibrillary tangles (*67*). p-TauS356 regulates cell growth and development (*68, 69*), is associated with lysosomal impairment, and contributes to tau stabilization when PAR-1/MARK activity is elevated (*70*).

One of the strengths of this study is that we demonstrated the effect of rapamycin on the detrimental effect of Tau^3E^ in a mouse model. In HT22 cells overexpressing Tau^3E^, increased p-mTOR and p-p70s6K levels and p-TauS214, T231, and S356 levels were detected. These findings are in line with those of an earlier study showing that mTOR signaling regulates tau phosphorylation in a Drosophila model and that mTOR activation can increase tau-mediated neurodegeneration(*71, 72*). Rapamycin decreases tau phosphorylation at Ser214 through the regulation of cAMP-dependent kinase activity and reduces cortical tau tangles by 80% in P301S-mutant tau transgenic mice and the level of sarkosyl insoluble tau in the forebrain by 70% (*73*). Previous studies in animal and Drosophila models have shown that excessive activation of the mTOR pathway leads to abnormal phosphorylation of tau, activation of cell cycle regulators and, eventually, apoptosis or even death, which can be reversed by rapamycin (*72, 74*). Downstream p70s6k of mTOR can directly phosphorylate the tau protein in neuroblastoma cells (SH-SY5Y) or hippocampal neurons (*75*). Several upstream signals, such as insulin, extracellular signal-regulated kinase 1/2, glycogen kinase-3β and protein phosphatase 2A, can interact with mTOR to induce tau hyperphosphorylation and the generation of Aβ plaques. Our previous studies revealed that mTOR hyperactivation can lead directly to tau hyperphosphorylation and aggregation. We found that rapamycin had a protective effect on neuronal function, oxidative stress, and mitochondrial homeostasis. Accumulating evidence has demonstrated that abnormal mTOR signaling in the brain affects mitochondrial function and oxidative stress (*76*). These findings are in line with those of earlier studies showing that rapamycin can improve cognitive dysfunction in a 3×Tg AD mouse AD animal model through intervention in the mTOR pathway (*32*). In our present study, we showed increased mTOR/P70S6K signaling in the hippocampi of mice overexpressing Tau^3E^. Rapamycin treatment reversed abnormal mTOR/P70S6K signaling and improved spatial learning in mice. Given the coincident abnormal tau accumulation and mTOR/P70S6K activation in damaged neurons in AD, we confirmed that tau phosphorylation at the S214, T231 and S356 sites (Tau^3E^) induced cell death and that rapamycin partially rescued neuronal death in mice.

Additionally, we found that Tau^3E^ led to an increase in the expression of mitochondrial fission-related p-Drp1 and a decrease in the expression of mitochondrial fusion-related proteins, resulting in mitochondrial damage. The overexpression of Tau^3E^ in C57BL/6BL mice activated mTOR, increased the generation of ROS, and induced lipid peroxidation and oxidative DNA damage. mTOR inhibition alleviated mitochondrial disease in a mouse model of Leigh syndrome (*77*). These findings are in line with an earlier study showing that an increased volume of mitochondria moving in neuronal axons reduces the number of mitochondria and the ability of tau to bind to microtubules in P301L tau-knockout neurons (*78*). An earlier study revealed that tau protein overexpression can inhibit mitochondrial function by reducing the activity of antioxidant enzymes and affecting ATP synthesis and synaptic function via localization to the mitochondrial membrane, leading to neuronal synaptic dysfunction in an animal model (*79*). The accumulation of full-length h-Tau can reduce the cell survival rate and cause neurodegeneration by regulating dynamic changes in mitochondrial fission and fusion in cells and animals (*80*). Human tau can result in the elongation of mitochondria as well as mitochondrial dysfunction and cell death by blocking the aggregation of Drp1 in mitochondria in Drosophila and mouse neurons (*81*). Hyperphosphorylated tau can directly interact with Drp1 to induce mitochondrial fission and excessive fragmentation in the AD brain (*82*).

Our study demonstrated that rapamycin treatment promoted comprehensive metabolic reprogramming across three integrated axes and restored the activity of intracellular calcium homeostasis and neurotransmitter homeostasis in the inosine-PPP-glycolysis axis. The spatial metabolic architecture of the hippocampus is profoundly disrupted by the accumulation of pathological tau. Overexpressing Tau^3E^ exhausted purine catabolites and glycolytic intermediates, which reflected a state of severe bioenergetic failure. Rapamycin treatment reversed these alterations, specifically by increasing the levels of inosine, uric acid, D-myo-inositol 1,4-bisphosphate and inositol signaling. Inosine has been increasingly recognized as a potent endogenous neuroprotectant that can serve as an alternative energy substrate under metabolic stress. Most of the previous studies have performed MALDI-MSI and other spectrometry investigation in models of amyloid-beta pathology (*83–86*). Recent study in PS19 tau mice showed similar changes in fructose 1,6-bisphosphate (m/z 338.9814) as in our study (*16*). Furthermore, the upregulated expression of PPP markers (D-ribose 5-phosphate and D-sedoheptulose 7-phosphate) suggested a redirection of glucose flux. Second, our data demonstrated that rapamycin rescues the structural lipidome, notably increasing long-chain polyunsaturated phospholipids such as PC (16:0/22:6). Restoring these DHA-enriched phosphatidylcholines is vital for preserving membrane fluidity and transmembrane protein activity. Ultimately, by replenishing essential phospholipids (PC, PE, and PI), rapamycin fortifies the neuronal membrane architecture, offering robust protection for hippocampal neurons against tau proteotoxicity. Third, rapamycin treatment led to the enrichment of Cer(d18:1/18:0) and SM(d16:1/20:5), which are vital for neuroplasticity and cellular resilience.

This study has several limitations. First, the rapamycin treatment (i.p.) used in the current study is rather high and of short duration. Both long-term and short-term rapamycin treatments had beneficial effects in earlier studies. Further analysis is needed to optimize the treatment plan. Second, we did not include female mice in the experiments. Previous studies have shown that the rapamycin-mediated increase in lifespan in mice is dose- and sex-dependent and is metabolically distinct from dietary restriction (*87*). Third, the downstream pathways influenced by rapamycin treatment were not investigated.

## 5 Conclusions

In summary, we demonstrated that p-Tau at S214, S356 and T231 increased mTOR expression, leading to the activation of ROS, mitochondrial and synaptic damage, and cognitive impairment. Rapamycin (i.p.) treatment reversed these tau-induced alterations including associated metabolic changes, and conferred neuroprotection in the Tau³LJ model. Rapamycin has potential therapeutic significance for tau-related ROS and mitochondrial dysfunction in AD and primary tauopathies.

## Authors’ Contributions

ZT and RN designed the experiments. MG, BL, YW and YTD carried out the data collection and collation, and XY, MG and ZT analyzed the data. ZZG and XLQ conducted the experiments and wrote the manuscript draft. ZT and RN revised the manuscript. All the authors reviewed the manuscript and approved the manuscript before submission.

## Data statement

The raw data are available upon request.

---

**Supplemental Table 1.** Antibodies used in this study.

| Antibody | Host | WB/IF<br>Dilution | Sources | Catalog No |
| --- | --- | --- | --- | --- |
| Anti-4-Hydroxynonenal | r | 1:200 | Abcam | #ab46545 |
| Anti-8-Hydroxy-2'-deoxyguanosine | m | 1:200 | Abcam | #ab62623 |
| Anti-beta-Actin | m | 1:15000 | affbiotech | #T0022 |
| Anti-mouse IgG (H+L) (HRP) | g | 1:5000 | Thermo Fisher | #31430 |
| Anti-Mouse IgG (H+L) Alexa Fluor™ 488 | d | 1:200 | Thermo Fisher | #A21202 |
| Anti-rabbit IgG (H+L) (HRP) | g | 1:5000 | Thermo Fisher | #31460 |
| Anti-Rabbit IgG (H+L) Alexa Fluor™ 546 | d | 1:200 | Thermo Fisher | #A10040 |
| Drp1 | r | 1:1000 | Abcam | #ab184247 |
| Fis1 | m | 1:500 | Santa | #sc-376447 |
| GAPDH | r | 1:6000 | Genetex | #GTX100118 |
| HT7 | r | 1:1000 | Invitrogen | #MN1000 |
| Mfn1 | m | 1:500 | Santa | #sc-166644 |
| Mfn2 | m | 1:500 | Santa | #sc-100560 |
| mTOR | r | 1:1000 | Cell Signaling Tech | #2983 |
| NeuN | r | 1:400 | Abcam | #ab177487 |
| NOX4 | r | 1:1000 | proteintech | #14347-1-AP |
| P70s6k | r | 1:1000 | Cell Signaling Tech | #2708 |
| pDrp1 | r | 1:1000 | Cell Signaling Tech | #3455s |
| pmTOR | r | 1 : 1000<br>/1:100 | Cell Signaling Tech | #5536s |
| pmTOR | m | 1:50 | Santa | #sc-293133 |
| pP70s6k | r | 1:1000 | Cell Signaling Tech | #9208 |
| pTau214 | r | 1:1000<br>/1:100 | Invitrogen | #44-742G |
| pTau231 | r | 1:5000<br>/1:500 | Abcam | #ab151559 |
| pTau356 | r | 1:1000<br>/1:100 | Abcam | #ab75603 |
| Tomm20 | r | 1:300 | Abcam | #ab186734 |
| β-tublin | r | 1:5000 | Beyotime | #21463 |

APP-CTFs, amyloid precursor protein-derived C-terminal fragments; ERK, extracellular signal-regulated kinase; GAPDH, glyceraldehyde 3-phosphate dehydrogenase; HRP, horseradish peroxidase; d: donkey; g: goat; P: phosphorylated; r: rabbit; m: mouse; T: total; dilution was used for western blot or immunofluorescence staining.

**Supplemental Table 2.** Chemicals and kits used in this study.

| Chemical/kit | Sources | Catalog No |
| --- | --- | --- |
| 2',7'-dichlorofluorescein diacetate assay kit | Beyotime, China | #S0033S |
| 5× FD TM DualColor Protein Loading Buffer | Fude Biological Technology, China | #FD006 |
| DAPI Fluoromount-G | Southern Biotech, USA | #0100-20 |
| Dimethyl sulfoxide | Sigma □ Aldrich | #D2650 |
| Dulbecco's modified Eagle's medium | Gibco, USA | #11960044 |
| Fetal Bovine Serum | Gibco, USA | #10099141C |
| Immobilon® Western Chemiluminescent HRP Substrate | Millipore, USA | #WBKLS0500 |
| Lipofectamine® 2000 Reagent | Invitrogen, USA | #11668-019 |
| Nonfat milk | Solarbio, USA | #D8340 |
| Phosphate-buffered saline | Biological Industries, Israel | #02-020-1A |
| Primary Antibody Dilution Buffer | Beyotime, China | #P0023A |
| PVDF membranes | Millipore, USA | #ISEQ00010 |
| Rapamycin (purity ≥98.0%) | MedChemExpress, , USA | #HY-10219 |
| Secondary Antibody Dilution Buffer | Beyotime, China | #P0023D |
| Trans-blot turbo 5x transfer buffer | Bio-Rad, USA | #10026938 |
| Triton™ X-100 | Sigma □ Aldrich | #T8200 |
| Tween-20 | Solarbio, USA | #T8220 |
| SDS polyacrylamide gels | Absin, China | #abs9382 |
| DAPI, 4',6-diamidino-2-phenylindole; HRP, horseradish peroxidase; PVDF, polyvinylidene fluoride; SDS, sodium dodecyl sulfate; |  |  |

**Supplemental Table 3.**
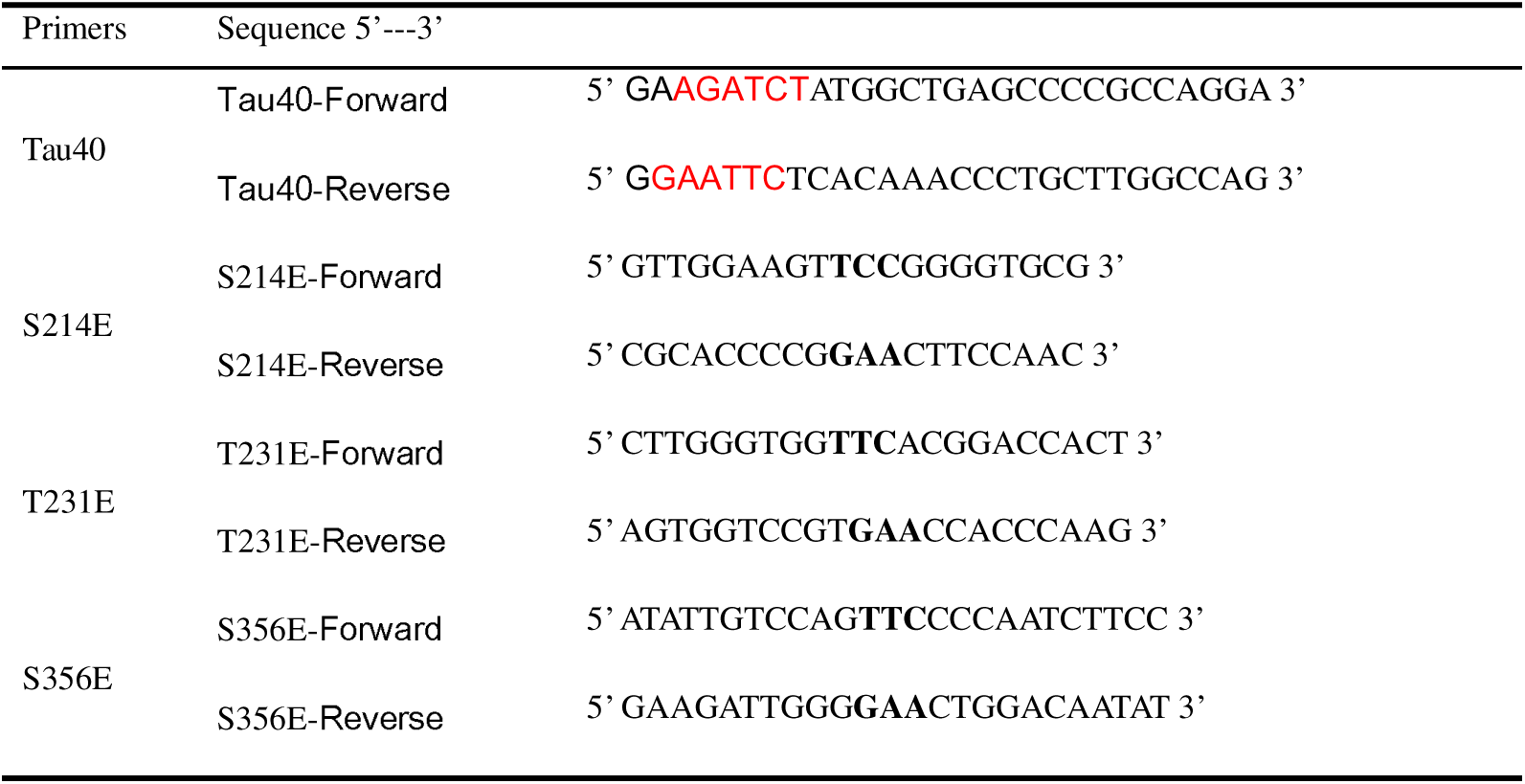
Designing the sequence of the phosphorylation point mutation primer.

**Supplemental Table 4.**
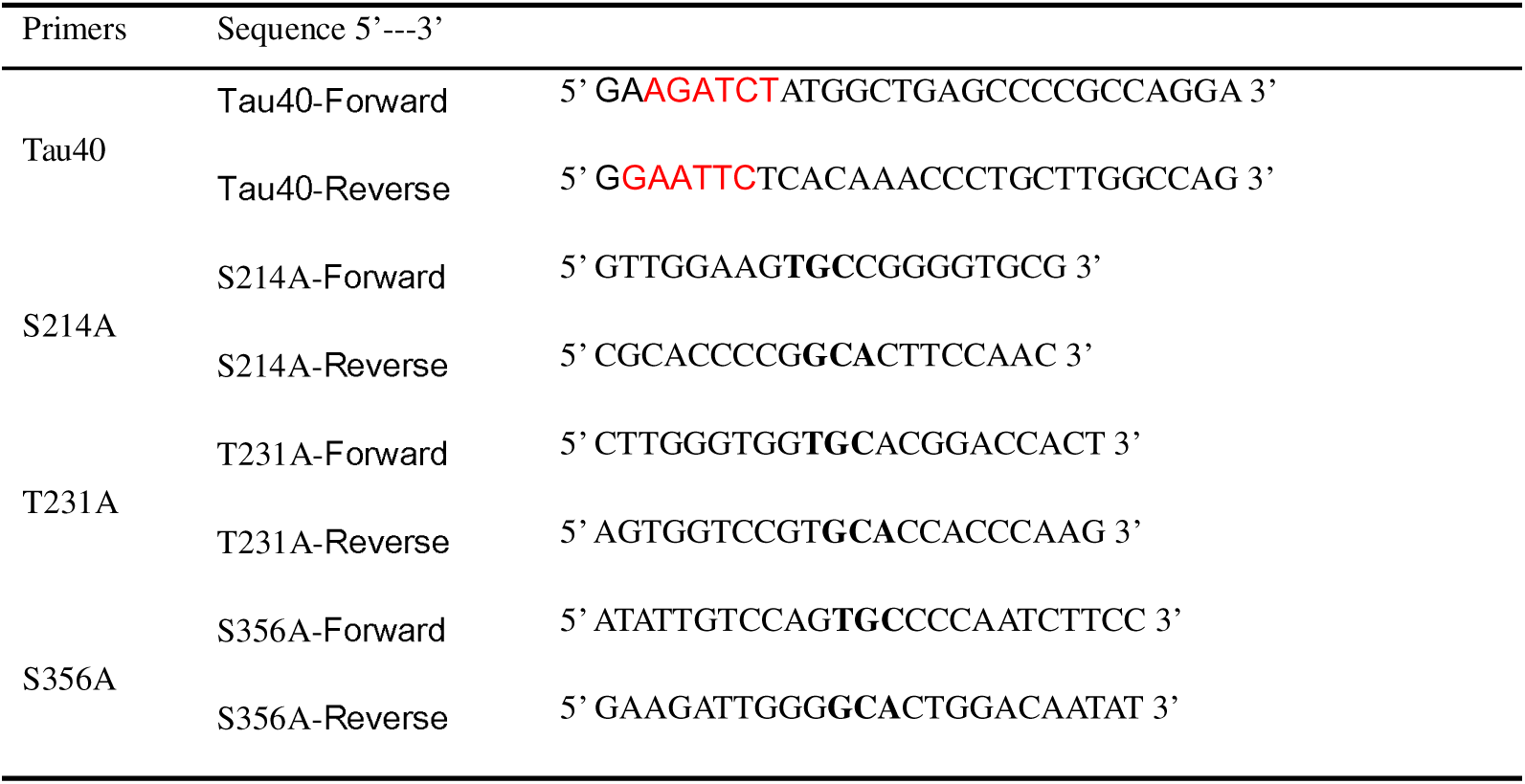
Dephosphorylation point mutation primer design sequence.

**Supplemental Table 5.** Details of the human brain samples used in this study.

| No | Diagnosis | Sex | Age | PMD (h) | ApoE | Amyloid | Braak stage |
| --- | --- | --- | --- | --- | --- | --- | --- |
| 1<sup>*#</sup> | AD | f | 76 | 5.08 | 44 | C | 5 |
| 2<sup>#&</sup> | AD | f | 80 | 5.33 | 44 | C | 6 |
| 3<sup>&</sup> | AD | m | 73 | 5.00 | 43 | C | 6 |
| 4<sup>*&</sup> | AD | f | 72 | 3.75 | 43 | C | 6 |
| 5<sup>*#</sup> | AD | f | 85 | 6.17 | 43 | C | 5 |
| mean[SD] |  | 4F/1 M | 77.2[13] | 5.07[2.42] | ApoE ε4 carrier<br>100% |  |  |

\*Tau231 staining; #Tau214 staining; &Tau356 staining; F, female; M, male; mean [SD].

